# Differential condensation of sister chromatids coordinates with Cdc6 to ensure distinct cell cycle progression in *Drosophila* male germline stem cell lineage

**DOI:** 10.1101/2021.03.08.434490

**Authors:** Rajesh Ranjan, Jonathan Snedeker, Matthew Wooten, Carolina Chu, Sabrina Bracero, Taylar Mouton, Xin Chen

**Affiliations:** Department of Biology, The Johns Hopkins University, Baltimore, MD 21218, USA; Fred Hutchinson Cancer Research Center, Seattle, WA 98109-1024, USA; Integrated Dermatology, Washington DC, 20036, USA; Department of Immunology, Harvard Medical School, Boston, MA 02115, USA

## Abstract

Stem cells undergo asymmetric division to produce both a self-renewing stem cell and a differentiating daughter cell. During *Drosophila* male germline stem cell (GSC) asymmetric division, preexisting old histones H3 and H4 are enriched in the self-renewed stem daughter cell, whereas the newly synthesized H3 and H4 are enriched in the differentiating daughter cell. However, the biological consequences in the two daughter cells resulting from asymmetric histone inheritance remained to be elucidated. In this work, we track both old and new histones throughout GSC cell cycle using high spatial and temporal resolution microscopy. We find several unique features differentiating old versus new histone-enriched sister chromatids, including nucleosome density, chromosomal condensation, and H3 Ser10 phosphorylation. These distinct chromosomal features lead to their differential association with Cdc6, an essential component of the pre-replication complex, which subsequently contributes to asynchronous initiation of DNA replication in the two resulting daughter cells. Disruption of asymmetric histone inheritance abolishes both differential Cdc6 association and asynchronous S-phase entry, demonstrating that asymmetric histone acts upstream of these critical events during cell cycle progression. Furthermore, GSC defects are detected under these conditions, indicating a connection between histone inheritance, cell cycle progression and cell fate decision. Together, these studies reveal that cell cycle remodeling as a crucial biological ‘readout’ of asymmetric histone inheritance, which precedes and could lead to other well-known readouts such as differential gene expression. This work also enhances our understanding of asymmetric histone inheritance and epigenetic regulation in other stem cells or asymmetrically dividing cells in multicellular organisms.

## Introduction

A fundamental question in developmental biology is how cells with identical genomes take distinct cell fates. One important context for understanding cell fate decision is ACD (Asymmetric Cell Division), wherein two daughter cells establish different cell fates following a single mitosis. ACD has been characterized in multiple organisms during development, homeostasis, and tissue regeneration (Blanpain and Fuchs, 2014; Sunchu and Cabernard, 2020; Venkei and Yamashita, 2018; Zion et al., 2020). Adult stem cells, in particular, often undergo ACD to generate one self-renewing stem daughter cell and another differentiating daughter cell.

Epigenetic mechanisms play important roles in cell fate decisions by altering chromatin structure and gene expression patterns while preserving primary DNA sequences (Allis and Jenuwein, 2016). In eukaryotic organisms, chromatin is organized by nucleosome units of 145-147-bp DNA wrapped around a histone octamer composed of H3, H4, H2A, and H2B (Kornberg, 1974; Luger et al., 1997; Richmond and Davey, 2003). Histones are one major epigenetic information carriers, serving both to package DNA and regulating gene expression through multiple mechanisms (Bannister and Kouzarides, 2011; Stillman, 2018; Zhang et al., 2021). In addition, the composition of nucleosome array and the positioning of individual nucleosomes play important roles in regulating local gene expression as well as DNA replication timing (Allshire and Madhani, 2018; Jiang and Pugh, 2009; Rhind and Gilbert, 2013; Struhl and Segal, 2013). For example, euchromatin with ‘open’ features often harbor less nucleosome density and active histone modifications at expressed genes and at early replication regions. In contrast, heterochromatin with ‘closed’ features accumulates with high nucleosome density and repressive histone modifications at silenced genes and at late replication regions. During cellular differentiation, both histone modifications and nucleosome density or positioning could change dramatically (Belsky et al., 2015; Goldberg et al., 2007; Lipford and Bell, 2001; Meshorer and Misteli, 2006; Rodriguez et al., 2017; Simpson, 1990; Stancheva, 2011). In multiple stem cell lineages, epigenetic mechanisms play critical roles including maintaining stem cell identity, specifying the differentiating cell fates, etc. (Atlasi and Stunnenberg, 2017; Avgustinova and Benitah, 2016; Feng and Chen, 2015b; Wooten et al., 2020b).

The *Drosophila* germline stem cell (GSC) systems provide great models to study adult stem cells (Fuller and Spradling, 2007; Vidaurre and Chen, 2021). Male GSCs divide asymmetrically to produce a self-renewing GSC and a differentiating gonialblast (GB) (Kiger et al., 2001; Leatherman and Dinardo, 2010; Tulina and Matunis, 2001; Yamashita et al., 2003). The GB undergoes transit-amplifying divisions as spermatogonial cells (SG) and then enters meiosis and terminally differentiation into mature sperm (Fuller, 1998; White-Cooper et al., 2000). Previously, we have shown that old H3 selectively segregates to the GSC whereas new H3 enriches in the GB during ACD of GSCs (Tran et al., 2012). Mis-regulation of asymmetric histone inheritance results in both GSC loss and progenitor germ cell tumors (Xie et al., 2015). The histone asymmetry is established during DNA replication *via* strand-specific incorporation and biased replication fork movement (Wooten et al., 2019). During ACD, epigenetically distinct sister chromatids are differentially recognized and segregated (Ranjan et al 2019) (Figure 1A). However, questions remain regarding how epigenetically distinct sister chromatids could regulate potentially distinct cellular and molecular features in the two resulting daughter cells.

**Figure 1.**
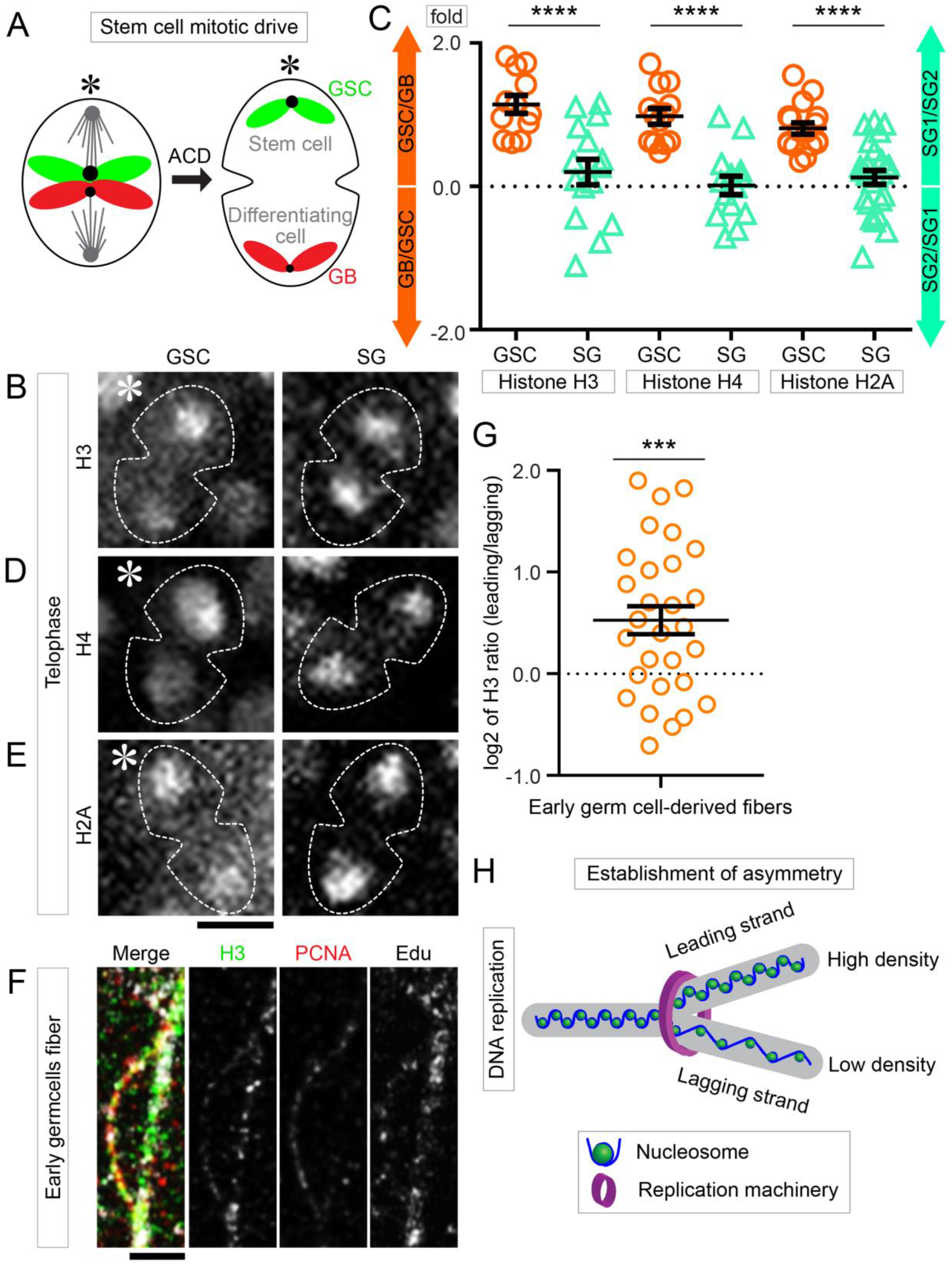
Global Nucleosome Density is Asymmetric during Male *Drosophila* GSC ACD. (**A**) Cartoon of the asymmetric segregation of old histone (green) and new histone (red) enriched sister chromatids in male *Drosophila melanogaster* GSCs. (**B**, **D**, **E**) Live images for total H3 (**B**), H4 (**D**), and H2A (**E**) for both GSCs (left column) and SGs (right column) in telophase. (**C**) Quantification of total histone (H3, H4, and H2A) segregation in telophase by live cell imaging for both GSCs and SGs, reflects nucleosome density (Table S1-3). Histone H3, GSC 1.50 ± 0.07 (n=12) and SG 1.10 ± 0.07 (n=15); histone H4, GSC 1.42 ± 0.05 (n=13) and SG 1.02 ± 0.05 (n=14); histone H2A, GSC 1.33 ± 0.04 (n=17) and SG 1.06 ± 0.04 (n=25) (****p<10^-4^ by Mann-Whitney t-test). (**F**) Chromatin fiber showing the distribution of total H3 during DNA replication. (**G**) Quantification of total H3 distribution from early germline derived chromatin fibers [GSC 1.44 ± 0.16 (n=29); (Table S4) (***P<10^-3^ by one sample t-test)]. (**H**) Cartoon shows that nucleosome density is asymmetric to the leading strand during DNA replication. Scale bars are 5 μm (**B, D-E**), 2 μm (**F**). Asterisk: hub. All ratios = average ± SE.

A critical cell type-specific feature is their unique cell cycle progression. During stem cell differentiation, dedifferentiation or trans-differentiation, cell fate changes and cell cycle remodeling are often coupled (Dalton, 2015; Guo et al., 2014; Hu et al., 2019; Soufi and Dalton, 2016). For example, elongated G1 phase length has been shown to contribute to the decision toward self-renewal rather than differentiation in mouse embryonic stem cells (Li et al., 2012; Li and Kirschner, 2014). On the other hand, S phase provides a window of time to sort out epigenetic information during replication-coupled nucleosome assembly (Serra-Cardona and Zhang, 2018; Snedeker et al., 2017; Stewart-Morgan et al., 2020; Zhang et al., 2020). However, initiation of S phase in multicellular organisms is not well understood, as this process relies more on chromatin features than the DNA sequences in higher eukaryotic cells, such as nucleosome density and histone modifications (Lipford and Bell, 2001; Patel et al., 2006; Simpson, 1990). It has been shown that nucleosome density around origins underlies the formation of the pre-replicative complex (pre-RC) and, hence, the timing and efficiency of origin firing (Lipford and Bell, 2001; McCune et al., 2008; McGuffee et al., 2013; Muller et al., 2014; Raghuraman et al., 1997; Simpson, 1990; Yabuki et al., 2002). Importantly, these changes are not established solely in G1 phase but rather from G2/M to G1 phase in many organisms. In the pre-RC, Cdc6 is an essential component that facilitates the loading of the replicative helicase MCM2-7 onto DNA, along with Orc1-6 and Cdt1, in preparation for active DNA replication (Bell and Dutta, 2002; Bleichert et al., 2017; Costa et al., 2013; Mendez and Stillman, 2003; Randell et al., 2006; Speck et al., 2005; Speck and Stillman, 2007). Up to date, most of the studies on these processes are based on *in vitro* studies, unicellular organisms such as yeast, or in cultured cells. It is much less known how these critical cell cycle dependent events contribute to cell fate decisions in multicellular organisms *in vivo*.

## Results

### Asymmetric nucleosome density between sister chromatids in GSCs

Chromatin structure affects binding of key factors that regulate subsequent processes such as transcription and replication (Jerkovic et al., 2020). To better understand how chromatin structure changes in asymmetrically dividing stem cells, we performed high spatial and temporal resolution three-dimensional (3D) live cell imaging using fly lines that express distinct *histone-eGFP* transgenes in early-stage germ cells (Figure S1A). When tracking histone inheritance patterns during ACD of GSCs, higher levels of histone H3 were found to associate with sister chromatids segregated toward the future GSC side compared to the future GB side (Figure 1B, Movie S1, Table S1). Quantification showed that ~1.5-fold more H3 is inherited by the GSC than the histones inherited by the GB side (Figures 1C, S1B). By contrast, such an asymmetric H3 levels was not detected in symmetrically dividing progenitor SGs at telophase (Figures 1B-C, S1C, Movie S2, Table S1). Furthermore, both H4 (Figure 1D, Movie S3, Table S2) and H2A (Figure 1E, Movie S5, Table S3) displayed a similar pattern, with ~1.4-fold more H4 and ~1.3 fold more H2A enriched on the sister chromatids that segregate to the GSC side in telophase GSCs (Figure 1C). Contrastingly, no such an asymmetry of either H4 (Movie S4) or H2A (Movie S6) was detected in SGs (Figure 1C). The differences among H2A, H3, and H4 likely reflect that both H2A and H3 have variants, such as H3.3 and H2A.v (Ferrand et al., 2020; Henikoff and Smith, 2015; Szenker et al., 2011).

Next, we utilized a SRCF (Super-Resolution imaging of Chromatin Fiber) method to directly visualize replicative DNA with associated histone proteins at single molecule resolution (Wooten et al., 2020a; Wooten et al., 2019). The chromatin fibers were extracted from early-stage germ cells expressing an *H3-EGFP* transgene using *nanos>H3-EGFP* fly testes. Active DNA replication regions were labeled by a short-pulse of a thymidine analog 5-ethynyl-2’-deoxyuridine (EdU) and imaged using high-spatial resolution Airyscan microscopy (Sivaguru et al., 2018). Consistent with the asymmetric histones between sister chromatids during ACD, replicating or newly replicated sister chromatid chromatin fibers displayed an asymmetric distribution of H3-EGFP (Figure 1F). Co-immunolabeling of a lagging strand-enriched protein PCNA (proliferating cell nuclear antigen) (Yu et al., 2014) identified that H3 is more enriched at the PCNA-less leading strand (Figure 1F). Comparable to the ratio between segregated sister chromatids during ACD of GSCs, ~1.4-fold higher nucleosome density was detected at the leading strand relative to the lagging strand on average (Figure 1G, Table S4). Notably, the current histone labeling strategy cannot distinguish pure GSC-derived chromatin fibers from other early-stage germ cell-derived ones, which could result in the relatively wide distribution of data points (Figure 1G).

Together, these results suggest that the GSC-specific asymmetric nucleosome densities between sister chromatids are likely established during S phase and segregated during M phase, and the sister chromatids carrying more nucleosomes are inherited by the self-renewing GSC (Figures 1H, S1D).

### Differential condensation of sister chromatids enriched with old versus new histones

Considering this new finding of nucleosome density difference between sister chromatids with the previous findings that old versus new histone enriched sister chromatids are established during DNA replication (Wooten et al., 2019) and recognized in mitosis (Ranjan et al., 2019), we hypothesize that the GSC-inherited sister chromatids are enriched with old histone and have higher nucleosome density. To visualize this, we performed high spatial resolution fixed cell imaging using a dual color system to label old H3 with eGFP and new H3 with mCherry (Figure S1A). In mid G2-phase GSCs, old and new H3 showed largely overlapping pattern (Figure S2A).

From late G2 phase, old and new H3 began to display distinct localization patterns (Figure S2B). After entering M phase, old H3-enriched regions condensed prior to new H3-enriched domains in prophase (Figures S2C-D). This differential condensation persisted from prophase to prometaphase and metaphase (Figures 2A-B, S2E-G). When old versus new histone-occupied chromatin regions were measured, a compaction factor was computed as the ratio of new H3-occupied area to old H3-occupied area (Figure S2H). This quantification showed on average a 2.64 compaction factor in prometaphase GSCs (Figure 2C, Table S5) whereas a 1.35 compaction factor in prometaphase SGs (Figures 2C-E, S2I-M, Table S5).

**Figure 2.**
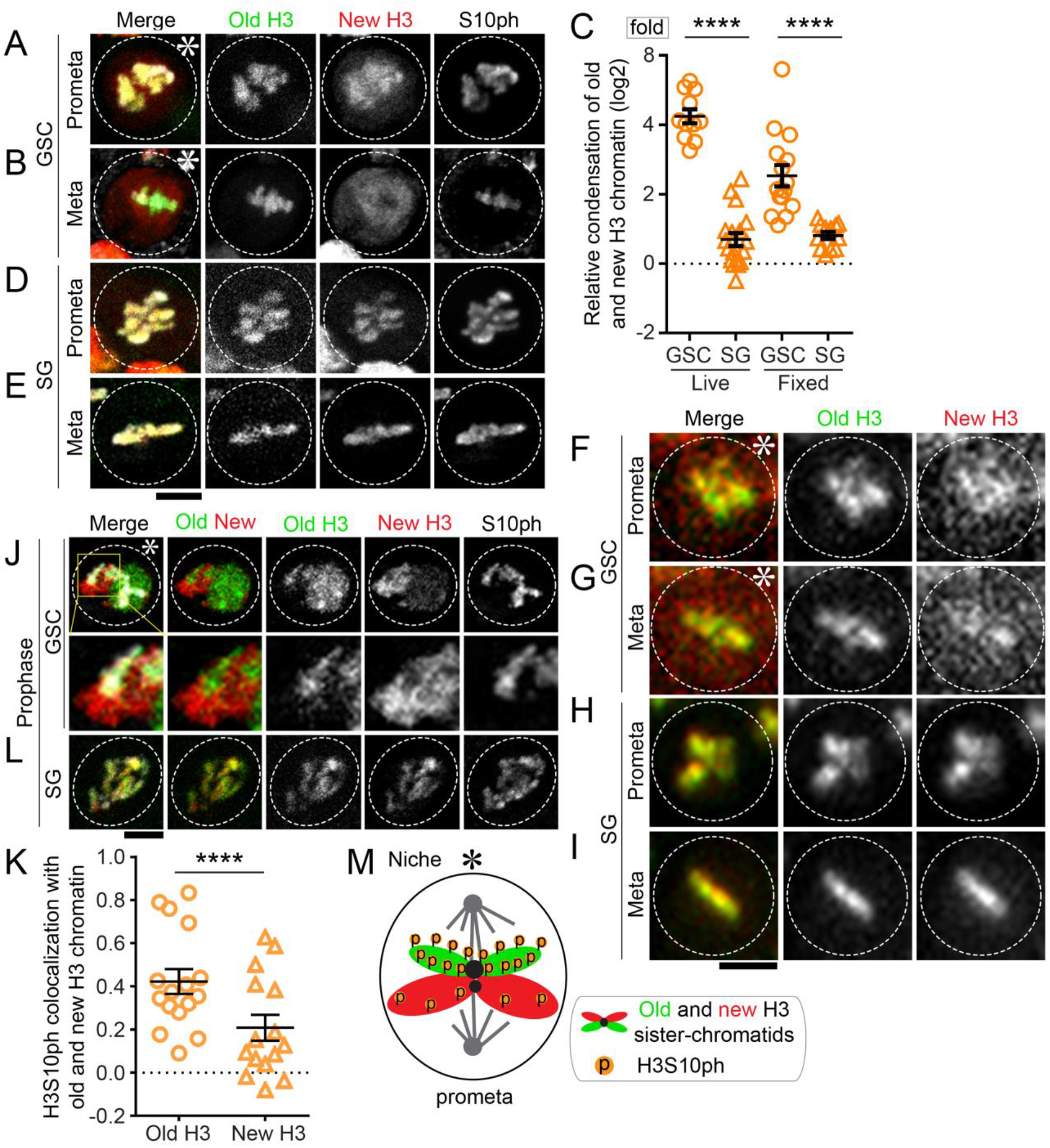
Old Histone Enriched Chromatin Condenses More and Earlier than New Histone Enriched Chromatin in Mitosis. (**A, B, D, E**) Fixed cell images of old and new H3 during prometaphase for GSCs (**A**), metaphase for GSCs (**B**), prometaphase for SGs (**D**), and metaphase for SGs (**E**). (**C**) Quantification of the relative condensation score for old vs. new histone chromatin in both GSCs and SGs using both live and fixed imaging [Live GSC 4.46 ± 0.32 and SG 1.32 ± 0.09; fixed GSC 2.64 ± 0.36 and SG 1.33 ± 0.05; (Table S5) (****p<10^-4^ by Mann-Whitney t-test)]. (**F-I**) Live cell image snapshots of prometaphase for GSCs (**F**), metaphase for GSCs (**G**), prometaphase for SGs (**H**), and metaphase for SGs (**I**). (**J, L**) Airyscan fixed cell images of the association of the histone modification H3S10ph with old rather than new histones in GSCs during asymmetric condensation in GSCs in prophase (**J**), but symmetric association in SGs (**L**). (**K**) Spearman colocalization of H3S10ph with old and new histones in GSCs [old H3 0.49 ± 0.05 and new H3 0.21 ± 0.06 (n=16); (Table S6) (****p<10^-4^ by Mann-Whitney t-test)]. (**M**) Cartoon of prometaphase asymmetric condensation of sister chromatids and preferential H3S10ph on old histone enriched sister chromatids. Scale bars are 2 μm. Asterisk: hub. All ratios = average ± SE.

To confirm these results using fixed cell imaging, we also performed high temporal resolution live cell imaging with the dual-color labeled old and new H3 (Figures S2N-S, Movie S7). Consistently, old and new H3 were largely overlapping in mid G2 phase (Figure S2N) but started to display distinct localization patterns in late G2 phase (Figure S2O). Further, distinct condensation patterns could be detected in early prophase (Figure S2P) and persisted through late prophase to metaphase (Figure 2C, 2F-G, S2P, Movie S7, Table S5). The average compaction factor was 4.46 in GSCs but 1.32 in SGs at late prophase to prometaphase (Figures 2C, 2H-I, S2S, Movie S8). This higher compaction factor in GSCs using live cell imaging than using fixed cell imaging could be due to more “free” new histones that are washed away in fixed cells. Together, these data demonstrate that old and new H3-enriched chromatin condense differentially in GSCs but not in SGs.

Old and new H3 are shown to carry distinct post-translational modifications by previous studies. For example, phosphorylation of many residues including both Ser10 and Thr3 of H3 (H3S10ph and H3T3ph) are enriched on old H3 in human cells using mass spectroscopy (Lin et al., 2016). H3S10ph is also critical for chromosome condensation in mitosis (Sawicka and Seiser, 2012). We found that H3S10ph preferentially co-localizes with old H3 in GSCs from early prophase to prometaphase (Figures 2J-K). By contrast, no such a preferential association of H3S10ph with old versus new H3 was detected in SGs (Figures 2K-L, Table S6). These data are consistent with the previous finding that the H3T3ph has more co-localization with old H3 than with new H3 in GSCs (Xie et al., 2015). In summary, these results indicate that old and new H3-enriched sister chromatids condense differentially in GSCs, likely due to their different phosphorylation at H3S10 (Figure 2M).

### Differential association of a key pre-RC component Cdc6 to sister chromatids in GSCs

These findings that sister chromatids with asymmetric nucleosome density condense differentially during ACD of GSCs raise the question of the biological consequences. As nucleosomes are found to compete with other DNA binding proteins (Ramachandran and Henikoff, 2016), we hypothesize that asymmetry of nucleosome density could make sister chromatids different substrates competing for other chromatin-associating proteins.

Cdc6 is an essential component of the pre-RC for initiating DNA replication in eukaryotic cells. Using CRISPR/Cas9 genome editing technology, we generated knock-in fly lines with a *mCherry* tag at the endogenous *cdc6* gene that encodes the Cdc6-mCherry fusion protein. We then performed high-spatial resolution fixed cell imaging to study the Cdc6 localization in mitotic early-stage germ cells. In GSCs, Cdc6 levels increased from late G2 to M phase but is largely excluded from the chromatin at prophase (Figures 3A, 3E). However, Cdc6 started to associate with chromatin in prometaphase and such an association increased at metaphase in GSCs (Figures 3B-E, S3A). By contrast, Cdc6 only associates with chromatin after exiting M phase and entering G1 phase in somatic cells (Paolinelli et al., 2009; Yim and Erikson, 2010). Intriguingly, in GSCs at telophase, Cdc6 displayed asymmetric association between segregated sister chromatids with enrichment toward the future GB side (Figure 3F). In contrast, such an asymmetry was not observed in SGs (Figure 3G).

**Figure 3.**
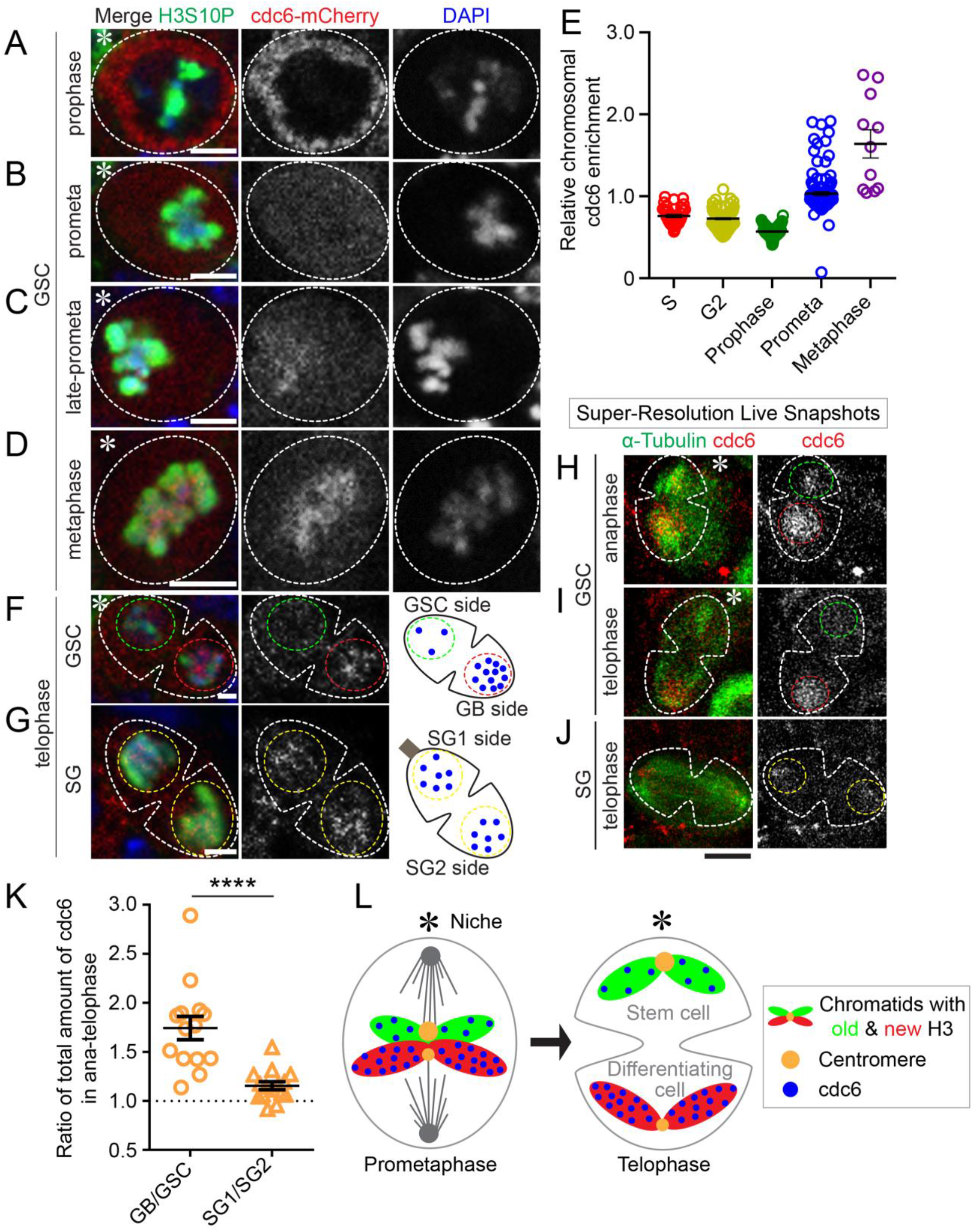
Cdc6 binds to chromatin during mitosis and segregate asymmetrically to the GB cell. (**A-D**) Cdc6 is initially localized in the cytoplasm during prophase (**A**) before gradually associating with the mitotic chromatin during prometaphase (**B, C**), and is strongly associated with chromatin by metaphase (**D**). (**E**) Quantification of the relative association of Cdc6 with the chromatin occupied region of the cell [S-phase 0.76 ± 0.02 (n= 36); G2-phase 0.73 ± 0.01 (n= 98); prophase 0.57 ± 0.01 (n= 70); prometaphase 1.03 ± 0.02 (n=120); metaphase 1.64 ± 0.17 (n =11)]. (**F, G**) Segregation of Cdc6 during mitosis in GSCs (**F**) and SGs (**G**) by fixed cell imaging. (**H-J**) Airyscan Super-Resolution Live Snaphshots (SRLS) for Cdc6 segregation during anaphase in GSCs (**H**), telophase in GSCs (**I**), and telophase in SGs (**J**). (**K**) Distribution of Cdc6 in anaphase/telophase cells for GBs relative to GSCs and SG pairs [GB/GSC 1.74 ± 0.12 (n=14); SG1/SG2 1.16 ± 0.04 (n=15); (Table S7)]. (**L**) Model for the differential binding of Cdc6 to new histone enriched sister chromatids before segregating preferentially to GBs. (****p<10^-4^ by Mann-Whitney t-test). Scale bars are 2 μm, (H-J) 5 μm. Asterisk: hub. All ratios = average ± SE.

To further validate the dynamics of Cdc6 in live cells, we performed SRLS (Super-Resolution Live Snapshots) imaging using Airyscan microscopy (Ranjan and Chen, 2021) to further examine mitotic germ cells with high spatial and temporal resolution (Figure S3B). Consistent with the fixed cell data, Cdc6 was excluded from the chromatin until prometaphase, when the levels of chromatin-associated Cdc6 increased (Figure S3C). In GSCs at anaphase and telophase, Cdc6 displayed asymmetry with higher levels associated with sister chromatids segregated toward the future GB (Figures 3H-I). However, in SG at telophase, such an asymmetric Cdc6 association was not detected (Figure 3J). Quantification of the SRLS imaging results showed that ~1.8-fold more Cdc6 associated with sister chromatids segregated towards the future GB side compared to sister chromatids segregated towards the future GSC side during ACD of GSCs (Figure 3K, Table S7). Contrastingly, a nearly equal level of Cdc6 was found between sister chromatids segregated in symmetrically dividing SGs (Figure 3K, Table S7). Collectively, these results demonstrate that Cdc6 differentially binds to sister chromatids and is asymmetrically inherited during ACD in GSCs (Figure 3L).

### Asynchronous initiation of DNA replication in post-mitotic GSC and GB nuclei

Given the essential role of Cdc6 in regulating origin licensing and S-phase initiation (Hossain and Stillman, 2016), we next sought to understand the impacts of asymmetric Cdc6 segregation on S-phase progression in the resulting GSC and GB after ACD. We first investigated DNA replication initiation by imaging the replication machinery component PCNA. In GSC-GB pairs derived from ACD, higher PCNA signal was detected in GBs compared to GSCs, indicating that GBs initiate DNA replication prior to GSCs (Figure 4A). Because following mitosis exit, condensed chromosomes decondense while S phase progresses, the size of nuclei could be used as a metric to represent relative extent of progression into S phase. Quantification of the PCNA ratio in GB nuclei to the daughter GSC nuclei revealed that pairs with small nuclear sizes, indicative of very early S phase, predominantly displayed more abundant PCNA in the GB nuclei compared to the GSC nuclei (Figure 4B, Table S8). With the progression of S-phase in both GBs and GSCs, judged by their larger nuclear sizes, the overall PCNA levels became more comparable (Figure 4B), consistent with previous reports that bulk DNA synthesis is simultaneously detected in GSC-GB pairs (Sheng and Matunis, 2011; Yadlapalli et al., 2011; Yadlapalli and Yamashita, 2013). Comparatively, such an asymmetry of PCNA was not detected in the symmetrically dividing SGs using the similar assays and criteria (Figures 4B-C).

**Figure 4.**
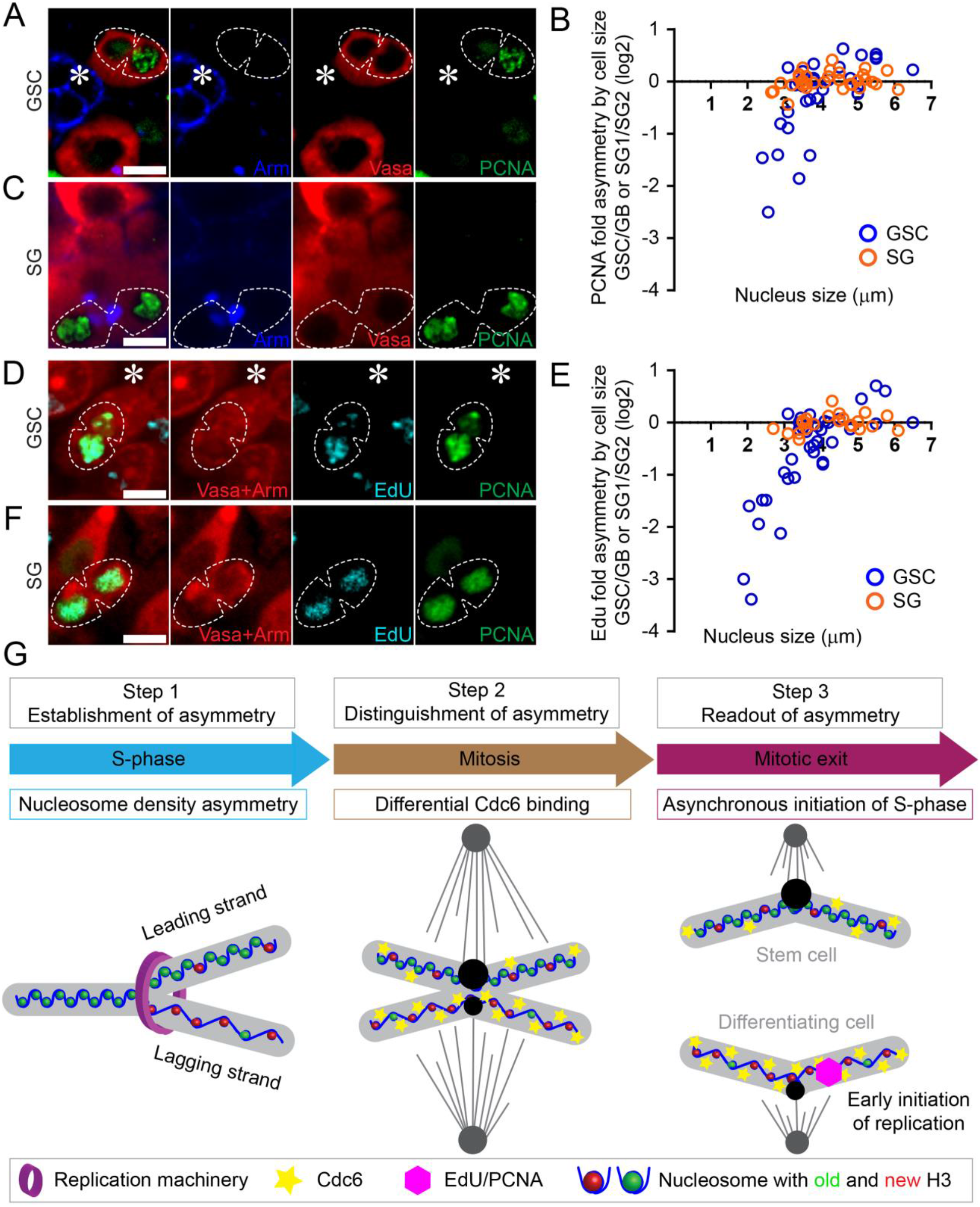
DNA Replication Begins Asynchronously Following GSCs Asymmetric Cell Division. (**A, C, D, F**) PCNA and EdU are enriched in GBs following division using fixed cell imaging. GBs are enriched for PCNA (**A**) in early post-mitotic pairs whereas SGs have equal levels of PCNA (**C**). (**B**) Quantification of PCNA asymmetry in GSCs (blue) and SGs (orange) using nuclear size as an indicator of time after division (n=33; Table S8). EdU as well as PCNA shows a similar enrichment in GBs (**D**), but not in SGs (**F**). (**E**) Quantification of EdU asymmetry in GSCs (blue) and SGs (orange) using nuclear size as an indicator of time after division (n=37; Table S9). (**G**) A model for how asymmetric histone inheritance could be upstream of the asymmetric entry into the next cell cycle via differential binding of Cdc6 to mitotic sister chromatids. Scale bars are 2 μm. Asterisk: hub.

To verify that the PCNA asymmetry reflects different timing of S phase initiation, we used a short pulse (~10-minute) of EdU to mark active DNA synthesis in GSC-GB pairs. Consistent with the PCNA results, EdU also displayed an asymmetric incorporation pattern in early S-phase GSC and GB, as more EdU was detected in GB nuclei while low EdU signal was detected in GSC nuclei immediately after exiting ACD, judged by their small nuclear sizes (Figures 4D-E, Table S9). Again, such an asymmetry of EdU incorporation was not detected in SGs (Figures 4E-F, Table S9). Overall, these results indicate that the sister chromatids inherited by GSCs and GBs are programmed differentially in the previous cell cycle to ensure asymmetric initiation of DNA replication and, hence, asynchronous cell cycle progression from M phase to early S phase (Figures 4G, S4A). The GB has almost negligible G1 phase. Consistently, precise measurement of the cell cycle time from metaphase of the previous cell cycle to metaphase of the subsequent cell cycle, using live cell imaging, demonstrated that GSCs have a longer cell cycle lasting for 14.85 hours on average (n=9) than that for GBs at 12.32 hours on average (n=7).

### Randomized sister chromatid inheritance abrogates asynchronous replication initiation

Previously, we showed that the microtubule depolymerizing drug Nocodazole (NZ) disrupts the temporal asymmetry of microtubules and results in randomized sister chromatid segregation in GSCs (Ranjan et al., 2019). Here, we utilized a similar strategy with NZ treatment followed by washout to acutely depolymerize microtubules (Figure 5A). We reason that if asynchronous initiation of DNA replication in post-mitotic GSC and GB nuclei is due to the distinct set of sister chromatids each nucleus inherits, this NZ-induced sister chromatid randomization should have impact on this asynchrony.

**Figure 5.**
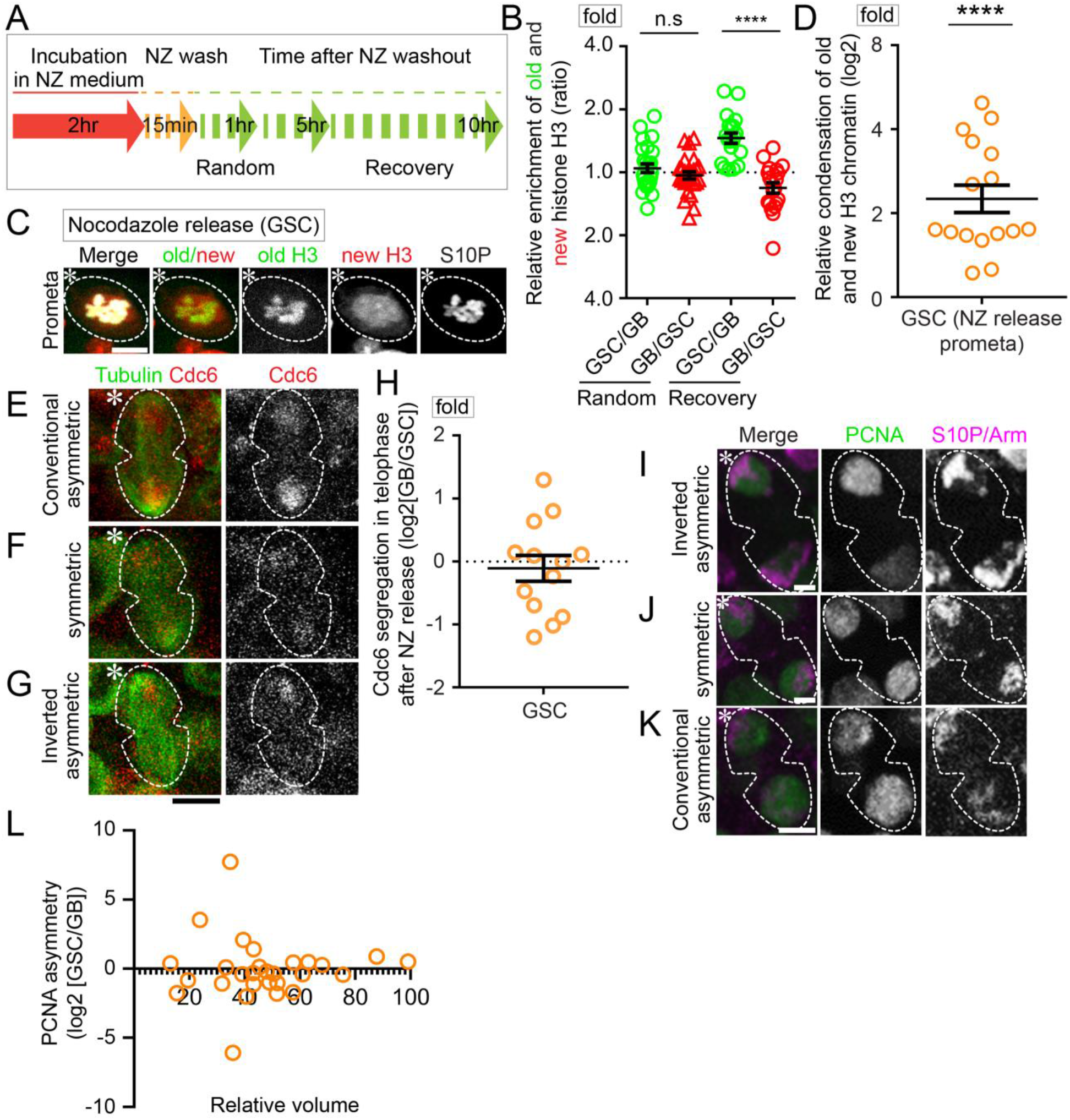
Nocodazole Randomizes the Relative Entry into S-Phase Timing by Randomizing Histone Segregation. (**A**) Schematic of Nocodazole (NZ) treatment and recovery experiments. (**B**) Quantification of Histone segregation immediately following release from NZ [old H3 GSC/GB = 1.08 ± 0.06; new H3 GB/GSC = 0.99 ± 0.04 (n=26); Table S10] and after recovering from NZ [old H3 GSC/GB = 1.51 ± 0.09; new H3 GB/GSC = 0.87 ± 0.05 (n=20); Table S10]. (**C**) Image of old and new histone differential condensation in GSC following NZ release, which is comparable to the pattern in the control GSCs without NZ treatment. (**D**) Relative old and new histone compaction is retained during Nocodazole (NZ) treatment as evidenced by an asymmetric compaction factor [compaction = 2.49 ± 0.30 (n=16); Table S11; (****p<10^-4^ by one sample t-test)]. (**E-G**) Airyscan Super-Resolution Live Snapshots show randomized Cdc6 segregation with Cdc6 preferentially segregating to the GB (E), symmetrically (**F**), or to the GSC (**G**). (**H**) Quantification shows that Cdc6 segregation patterns are random [GB/GSC = 1.05 ± 0.16 (n=13); Table S12]. (**I-K**) Fixed images of PCNA show that replication initiation timing is also randomized in early post-mitotic pairs as PCNA can segregate to the GSC (**I**), symmetrically (**J**), or to the GB (**K**). (**L**) Quantification of PCNA asymmetry in post-mitotic pairs using the relative nuclear volume as an approximation for time after division (n=28; Table S13). Scale bars are 2 μm (**C, I-K**), 5 μm (**E-G**). Asterisk: hub. All ratios = average ± SE.

First, to ensure that the NZ treatment is acute rather than persistent, a potential concern for secondary effects, we explored whether NZ-induced randomized sister chromatid segregation could be temporally controlled. Using a NZ treatment followed by washout regime (Figure 5A) and live cell imaging, we found mostly randomized old and new H3 inheritance patterns immediately after NZ release (Figures 5B, S5A), consistent with previous results (Ranjan et al., 2019). However, with a recovery after washing out NZ, the asymmetric old and new H3 inheritance patterns could be restored, with both displayed significant asymmetric patterns (Figures 5B, S5B, Table S10). These results demonstrate that NZ temporally disrupts asymmetric histone inheritance, but this effect could be reversed over time.

Next, we explored whether NZ treatment disrupts differential sister chromatid condensation (Figure 2), asymmetric Cdc6 inheritance (Figure 3), and the asynchronous initiation of DNA replication (Figure 4). Immediately after NZ release, old and new H3-enriched chromatin still displayed differential condensation at prometaphase, wherein old H3-enriched regions condensed 2.49-fold more than new H3-enriched regions in GSCs (Figures 5C-D, Table S11), comparable to the results in untreated GSCs (Figures 2C, S5C). These results indicate that the NZ treatment does not interfere with differential condensation of old and new H3-enriched chromatin, consistent with the hypothesis that the asymmetric old and new histones are established prior to mitosis and recognized by microtubules during mitosis, and only this latter step is disrupted by NZ (Ranjan et al., 2019). However, if Cdc6 preferentially binds to the less condensed new H3-enriched sister chromatids, then NZ treatment should randomize the Cdc6 segregation pattern, along with the randomized histone segregation pattern (Figure S5D). Indeed, live cell imaging results showed randomized Cdc6 segregation in anaphase and telophase GSCs immediately after NZ release, with asymmetric, symmetric, and inverted-asymmetric Cdc6 segregation (Figures 5E-H, Table S12). To further test whether asynchronous initiation of DNA replication changes, we studied the PCNA patterns in in the GSC and GB nuclei using fixed cell imaging. We found that NZ treatment also resulted in a randomized PCNA distribution with asymmetric, symmetric, and inverted-asymmetric PCNA patterns (Figures 5I-L, Table S13). Collectively, these results demonstrate that NZ treatment randomizes old and the new H3 inheritance, which results in randomized Cdc6 inheritance and asynchronous S-phase entry of the post-mitotic GSC and GB.

### H3S10A abolishes differential sister chromatid condensation and results in symmetric histone inheritance

Since the H3S10ph preferentially co-localizes with old H3 in GSCs (Figures 2J-K, 2M) and this modification is critical for chromosome condensation in mitosis, we speculated that H3S10ph could regulate the differential condensation between old and new H3-enriched sister chromatids in GSCs. To examine this possibility, we generated transgenic fly lines expressing a mutant H3S10A, in which the Ser10 residue was mutated to an unphosphorylable Alanine (Ala or A). When expressing the mutant H3S10A in early-stage germ cells including GSCs, the H3S10ph signals reduced to ~52.88% of the levels in the control cells (Figures S6A-B). The differential condensation between old and new H3S10A diminished with more overlapping patterns in GSCs from late G2 phase (Figure S6C) to M phase (prophase to metaphase in Figures S6E, 6A-D). Compared to the 2.64 compaction factor in H3-expressing GSCs, the compaction factor reduced to 1.36 in H3S10A-expressing GSCs (Figure 6E, Table S14), which is comparable of the 1.35 compaction factor in H3-expressing SGs (Figure 2C). Even though some differential condensation could still be detected in H3S10A-expressing GSCs (Figures S6D, S6F), there is a significant decrease of GSCs with differential condensation from 94% in H3-expressing GSCs to 38% in H3S10A-expressing GSCs (Figure 6F). Together, these data support the hypotheses that phosphorylation at H3S10 contributes to the differential sister chromatid condensation enriched with old versus new H3 in GSCs (Figure 2M) and expressing a mutant H3S10A greatly compromises differential sister chromatid condensation.

**Figure 6.**
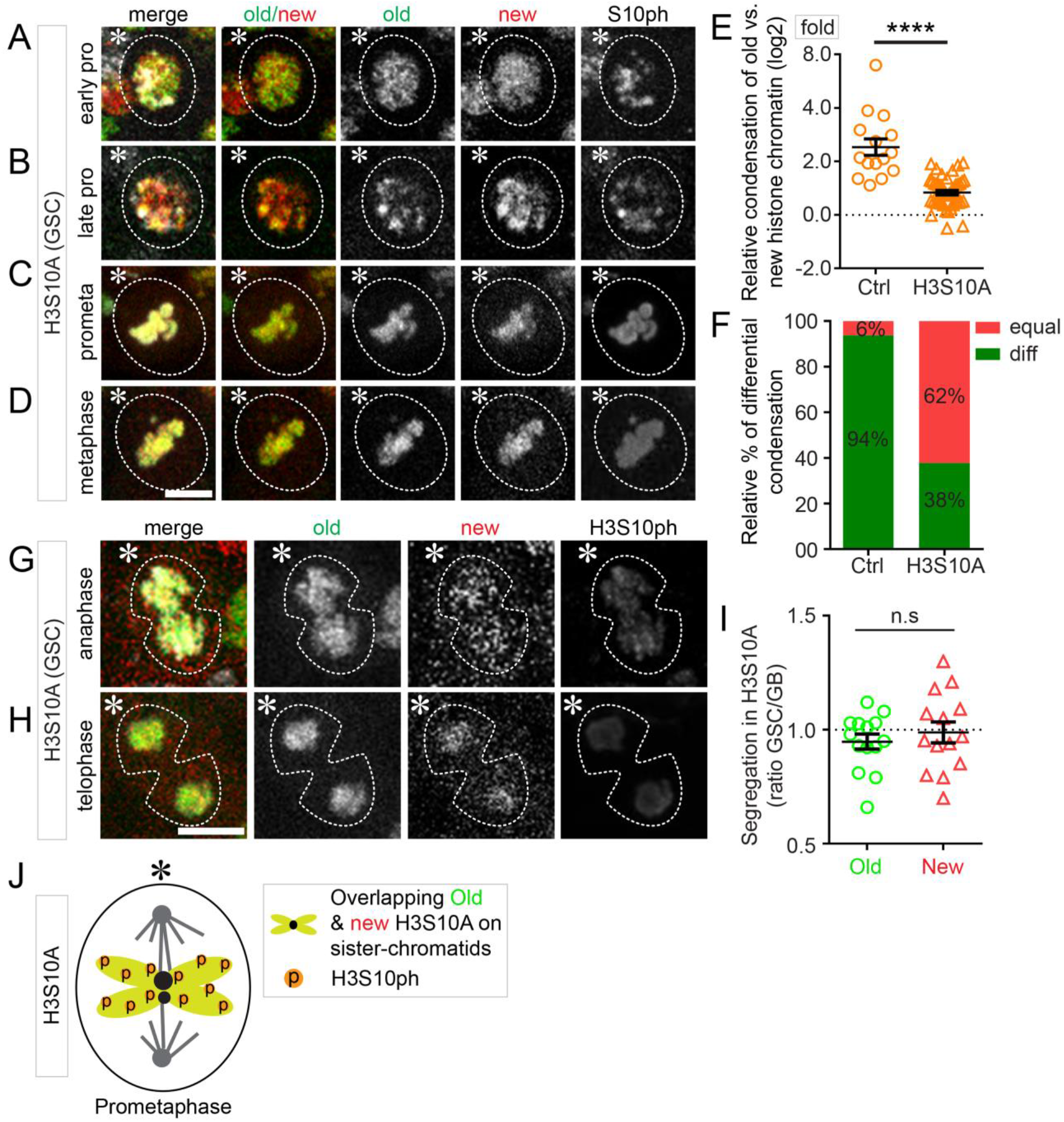
The H3S10A Mutation Eliminates Differential Mitotic Condensation and Asymmetric Histone Inheritance. (**A-D**) Old and new histone enriched chromatin condense equally in early prophase (**A**), late prophase (**B**), prometaphase (**C**), and metaphase (**D**) with the H3S10A mutant background. (**E**) Quantification of the relative condensation of old and new histone enriched chromatin in the control H3- and H3S10A-expressing GSCs [H3S10A compaction = 1.36 ± 0.04 (n=45); Table S14]. (**F**) The fraction of equally and differentially condensed mitotic sister chromatids in the control H3- and H3S10A-expressing GSCs. (**G-H**) Old and new histones segregate symmetrically in anaphase and telophase with the H3S10A mutant. (**I**) Quantification of the old and new histone segregation pattern following GSC division [old H3 = 0.95 ± 0.03; new H3 = 0.99 ± 0.05 (n=14); Table S15]. (**J**) Model of the H3S10A mutant resulting in equal H3S10 phosphorylation and equal sister-chromatids condensation. Scale bars are 2 μm (**A-D**) 5 μm (**H-I**). Asterisk: hub. All ratios = average ± SE.

We next examined whether H3S10A affects asymmetric histone segregation pattern. Indeed, this mutant histone leads to mostly symmetric old and new H3S10A segregation in anaphase and telophase GSCs (Figures 6G-H). Quantification confirmed that both old H3S10A and new H3S10A displayed nearly symmetric inheritance patterns (Figure 6I, Table S15). Notable, these results are different from the NZ-induced randomized patterns occurring in mitosis, where asymmetric, symmetric, and inverted-asymmetric patterns were all detectable (Ranjan et al., 2019), suggesting that this mutant histone likely abolishes the establishment of histone asymmetry prior to mitosis. Collectively, these results indicate that the differential phosphorylation of old and new H3 at Ser10 is required for asymmetric H3 inheritance in GSCs and expressing a mutant H3S10A results in a symmetric histone inheritance pattern (Figure 6J).

### H3S10A changes asynchronous initiation of DNA replication in post-mitotic GSC-GB pairs and leads to GSC defects

Next, we explored whether symmetric histone inheritance resulted from H3S10A expression also leads to changes of the asynchronous S-phase entry of the post-mitotic GSC-GB pairs, using the EdU-pulse labeling experiment (Figures 4D-F). When expressing H3S10A, GSC and GB enter the subsequent early S phase synchronously, shown by the nearly equal incorporated EdU levels and the similar DNA replication initiation patterns in both nuclei (Figure 7A). Further analyses of the EdU levels in early S-phase GSC-GB pairs, judged by their small nuclear size, revealed an almost equal amount of EdU in both nuclei (Figure 7B, Table S16). Together, these results suggest a nearly synchronous initiation of DNA replication in both H3S10A-expressing GSC and GB nuclei.

**Figure 7.**
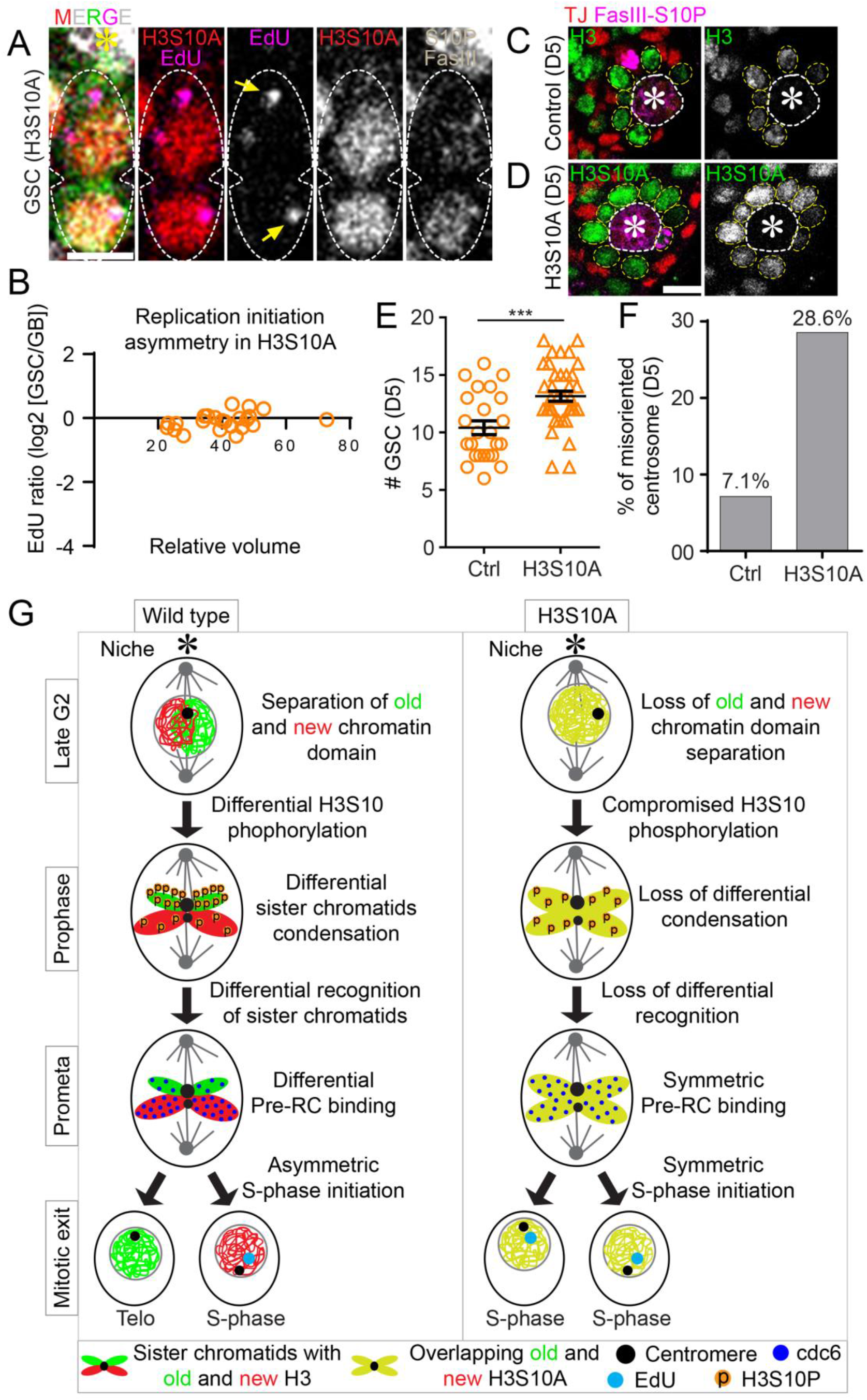
H3S10A Induces Symmetric Entry into the Next Cell Cycle. (**A**) EdU levels are comparable in a late telophase GSC division. (**B**) Quantification of EdU distribution in postmitotic pairs following GSC division shows that the two cells enter the next cell cycle synchronously (n=25, Table S16). (**C**, **D**) Images showing an expanded number of GSCs five days after eclosion in the H3S10A-expressing (**D**) compared to the control H3-expressing testes (**C**). (**E**) Quantification of the number of GSCs five days after eclosion in the control H3- and the H3S10A-expressing testes [Ctrl, Average = 10.41 (n=29); H3S10A, Average = 13.16 (n=38)]. (**F**) Quantification of the misoriented centrosome five days after eclosion in the control H3-expressing and the H3S10A-expressing GSCs [Ctrl = 7.13% (n=463); H3S10A = 28.57% (n=329)]. (**G**) Model for the role of differential condensation in driving the asymmetric entry into the next S-phase in the control wild-type GSCs, but not in the H3S10A mutant. Scale bars are 5 μm. Asterisk: hub. All ratios = average ± SE.

Furthermore, we investigated whether H3S10A expression could result in any germ cell defects. Because H3S10ph is required for chromosomal condensation and other important cellular events such as transcription (Ivaldi et al., 2007; Kalitsis et al., 2017; Nowak and Corces, 2004; Sawicka and Seiser, 2012), we sought to avoid potential secondary defects by compromising H3S10 phosphorylation in early-stage germline only in adult flies. To achieve this, we turned on H3S10A expression in a spatiotemporally specific manner by pairing the temperature-sensitive Gal80 controlled by *tubulin* promoter (*tub-Gal80^ts^*) with the early germline-specific *nos-Gal4* driver (*tub-Gal80^ts^; nos-Gal4>H3S10A-GFP*). After shifting from permissive temperature to restrictive temperature to inactivate Gal80 and turn on H3S10A expression for five days (D5), there was a decrease of the mitotic index in H3S10A-expressing GSCs (1.52%, n=329), compared to H3-expressing *Ctrl* GSCs (2.38%, n=463). However, the average GSC number judged by their anatomic position adjacent to the hub structure had an increase (Figures 7C-E). Interestingly, a much higher percentage of the H3S10A-expressing GSCs carried misoriented centrosomes compared to the control GSCs (Figure 7F), suggesting increased dedifferentiated GSCs that are arrested from entering M phase due to the “centrosome orientation checkpoint” (Cheng et al., 2008; Venkei and Yamashita, 2015; Yamashita et al., 2007). Consistently, prolonged expression of H3S10A for 10 days (D10) led to decreased GSC number (Figures S7A-B), likely due to the eventually defective cell cycle progression. Finally, increased hub area was detected in H3S10A-expressing testes at D10 (Figure S7C), consistent with previous reports as a secondary defect due to GSC loss (Dinardo et al., 2011; Gonczy and DiNardo, 1996; Monk et al., 2010; Ranjan et al., 2019; Tarayrah et al., 2015; Tazuke et al., 2002; Xie et al., 2015).

Summarizing the results by expressing the H3S10A mutant in early-stage germline (Figure 6 and 7), we propose that the differential condensation of old and new H3 enriched sister chromatids is required for both asymmetric H3 segregation in GSCs and asynchronous DNA replication initiation in post-mitotic GSC and GB nuclei (Figure 7G). The cellular defects detected in H3S10A-expressing GSCs suggest that these coordinated chromosomal changes and regulated cell cycle progression are required for stem cell maintenance and proper activity.

## Discussion

Here, using a series of high spatiotemporal microscopy methods and molecular genetics approaches, we demonstrated multiple distinct features displayed by old *versus* new H3 enriched chromatin regions, including nucleosome density asymmetry, differences in the degree and timing of chromosome condensation, and differential phosphorylation of a critical H3 residue. Furthermore, we identified several key features that are likely ‘read-outs’ of histone asymmetry, such as differential binding of a crucial replication component, Cdc6, in a cell cycle dependent manner, and asynchronous initiation of DNA replication between the two daughter cells. Finally, disruption of these chromatin asymmetry leads to mis-regulation of the cell cycle and misbehavior of the stem cells. Together, these results suggest that there is an intimate relationship between chromatin structure, cell cycle progression and cell fate determination or maintenance.

The discovery of asymmetric histone inheritance during ACD of *Drosophila* male GSCs provides an *in vivo* system to study the impact of distinct epigenomes on cell fate determination at single-cell level in real-time. Here, our results provide evidence for a novel cell cycle remodeling process regulated by asymmetric histones. The distinct chromatin features between sister chromatids differentially regulate the subsequent cell cycle progression in the two resulting daughter cells. The nucleosome-dense, old H3-enriched sister chromatids are retained in GSCs whereas the nucleosome-sparse, new H3-enriched sister chromatids are inherited by the GB. Nucleosome occupancy and positioning are critical factors in regulating both transcription and replication. For example, nucleosome depleted regions (NDRs) specify promoters, enhancer, and replication origins (Lai and Pugh, 2017; Radman-Livaja and Rando, 2010). Actively transcribed regions as well as early replicating regions are often less occupied by nucleosomes. In addition, nucleosome occupancy and heterochromatin formation are also dynamically regulated in cellular differentiation during development. Typically, more pluripotent or progenitor cells tend to have more open chromatin structure with less nucleosome density while heterochromatin with dense nucleosomes increase with cellular differentiation (Chen and Dent, 2014; Yadav et al., 2018). Thus, it is interesting that in the *Drosophila* male GSC lineage, our results demonstrate lower nucleosome density in the more differentiating GB than the GSC, raising an intriguing possibility that this feature is related to the unique reprogramming processes in the germline. Distinct from cellular differentiation in somatic lineages, germline cellular differentiation is not toward a “terminal” end, but to be reset to start a new life cycle (Cinalli et al., 2008; Eun et al., 2010; Feng and Chen, 2015a).

Further, the new findings that sister chromatids with different nucleosome densities differentially associate with the replication initiation factor Cdc6 are consistent with previous findings that nucleosome density around origins underlies the formation of the pre-RC and origin firing (Lipford and Bell, 2001; McCune et al., 2008; McGuffee et al., 2013; Muller et al., 2014; Raghuraman et al., 1997; Simpson, 1990; Yabuki et al., 2002). While Cdc6 typically binds to DNA in G1 phase in somatic cells (Paolinelli et al., 2009; Yim and Erikson, 2010), the association of Cdc6 in early-stage *Drosophila* male germline appears to start in early to mid-M phase, likely in preparation for the immediate initiation of DNA replication in the GB nucleus without detectable G1 phase. Asymmetries in Cdc6 association could promote early S-phase entry in several ways. First, Cdc6 has been well characterized for its role in loading the MCM2-7 (Cook et al., 2002; Tanaka et al., 1997; Yuan et al., 2020). Therefore, higher levels of Cdc6 in GB chromatin could accelerate the loading of helicase proteins necessary for the initiation and progression of DNA replication. Second, Cdc6 has been found to directly activate *Cyclin E* transcription, an event in the progression from G1 to S phase (Hossain and Stillman, 2016).

Therefore, asymmetries in Cdc6 could help in the direct upregulation of a key cell cycle regulator necessary to drive GB cells into the next S-phase faster with almost a negligible G1 phase. The cascade of events described here reveals a surprising role for nucleosome density differences in helping remodel cell cycle programs following stem cell ACD to establish distinct cell fates.

In summary, our results demonstrate that the chromatin status differences between sister chromatid contribute to a series of cellular events that result in distinct remodeling of the subsequent cell cycle. As cell cycle progression and timing are inherent properties of each cell type, this work reveals that remodeling cell cycle could serve as a critical step for cellular differentiation in stem cell lineages. Up to date, understanding how epigenetic mechanisms regulate cell identities mainly focus on their roles in differential gene expression. This work connects histone inheritance, cell cycle progression and cell fate decision, providing more crucial biological ‘readout’ of asymmetric histone inheritance. In the future, it will be interesting to examine how these features are related to the transcriptome changes, as dramatic transcriptome changes have been shown in the male GSC lineage (Shi et al., 2020). Moreover, the high spatial and temporal resolution microscopy methods developed in this work provides new opportunities to study stem cell properties in the context of continuous cell cycle at single-cell resolution *in vivo*, which could be widely applied to other stem cell lineages or asymmetrically dividing cells in multicellular organisms.

## Acknowledgements

We thank Chen lab members for suggestions. We thank Johns Hopkins Integrated Imaging Center for confocal imaging. Supported by NIH 5T32GM007231 (J.S., M.W.), F31 GM115149-01A1 (M.W.), NIH R35 GM127075 and R01 HD102474, and the Howard Hughes Medical Institute (X.C.)

## Author contributions

R.R., J.S., M.W. and X.C. conceptualized the study. R.R., J.S., M.W., C.C., S.B. performed all the experiments and most data analysis, T.M. helped with the data analysis. R.R., J.S., M.W. and X.C. wrote the manuscript.

## References

Allis, C.D., and Jenuwein, T. (2016). The molecular hallmarks of epigenetic control. Nat Rev Genet 17, 487–500.

Allshire, R.C., and Madhani, H.D. (2018). Ten principles of heterochromatin formation and function. Nat Rev Mol Cell Biol 19, 229–244.

Atlasi, Y., and Stunnenberg, H.G. (2017). The interplay of epigenetic marks during stem cell differentiation and development. Nat Rev Genet 18, 643–658.

Avgustinova, A., and Benitah, S.A. (2016). Epigenetic control of adult stem cell function. Nat Rev Mol Cell Biol 17, 643–658.

Bannister, A.J., and Kouzarides, T. (2011). Regulation of chromatin by histone modifications. Cell Res 21, 381–395.

Bell, S.P., and Dutta, A. (2002). DNA replication in eukaryotic cells. Annu Rev Biochem 71, 333–374.

Belsky, J.A., MacAlpine, H.K., Lubelsky, Y., Hartemink, A.J., and MacAlpine, D.M. (2015). Genome-wide chromatin footprinting reveals changes in replication origin architecture induced by pre-RC assembly. Genes Dev 29, 212–224.

Blanpain, C., and Fuchs, E. (2014). Stem cell plasticity. Plasticity of epithelial stem cells in tissue regeneration. Science 344, 1242281.

Bleichert, F., Botchan, M.R., and Berger, J.M. (2017). Mechanisms for initiating cellular DNA replication. Science 355.

Chen, T., and Dent, S.Y. (2014). Chromatin modifiers and remodellers: regulators of cellular differentiation. Nat Rev Genet 15, 93–106.

Cheng, J., Turkel, N., Hemati, N., Fuller, M.T., Hunt, A.J., and Yamashita, Y.M. (2008). Centrosome misorientation reduces stem cell division during ageing. Nature 456, 599–604.

Cinalli, R.M., Rangan, P., and Lehmann, R. (2008). Germ cells are forever. Cell 132, 559–562.

Cook, J.G., Park, C.H., Burke, T.W., Leone, G., DeGregori, J., Engel, A., and Nevins, J.R. (2002). Analysis of Cdc6 function in the assembly of mammalian prereplication complexes. Proc Natl Acad Sci U S A 99, 1347–1352.

Costa, A., Hood, I.V., and Berger, J.M. (2013). Mechanisms for initiating cellular DNA replication. Annu Rev Biochem 82, 25–54.

Dalton, S. (2015). Linking the Cell Cycle to Cell Fate Decisions. Trends Cell Biol 25, 592–600.

Dinardo, S., Okegbe, T., Wingert, L., Freilich, S., and Terry, N. (2011). lines and bowl affect the specification of cyst stem cells and niche cells in the Drosophila testis. Development 138, 1687–1696.

Eun, S.H., Gan, Q., and Chen, X. (2010). Epigenetic regulation of germ cell differentiation. Curr Opin Cell Biol 22, 737–743.

Feng, L., and Chen, X. (2015a). Epigenetic regulation of germ cells-remember or forget? Curr Opin Genet Dev 31, 20–27.

Feng, L., and Chen, X. (2015b). Epigenetic regulation of germ cells-remember or forget? Curr Opin Genet Dev 31, 20–27.

Ferrand, J., Rondinelli, B., and Polo, S.E. (2020). Histone Variants: Guardians of Genome Integrity. Cells 9.

Fuller, M.T. (1998). Genetic control of cell proliferation and differentiation in Drosophila spermatogenesis. Semin Cell Dev Biol 9, 433–444.

Fuller, M.T., and Spradling, A.C. (2007). Male and female Drosophila germline stem cells: two versions of immortality. Science 316, 402–404.

Goldberg, A.D., Allis, C.D., and Bernstein, E. (2007). Epigenetics: a landscape takes shape. Cell 128, 635–638.

Gonczy, P., and DiNardo, S. (1996). The germ line regulates somatic cyst cell proliferation and fate during Drosophila spermatogenesis. Development 122, 2437–2447.

Guo, S., Zi, X., Schulz, V.P., Cheng, J., Zhong, M., Koochaki, S.H., Megyola, C.M., Pan, X., Heydari, K., Weissman, S.M., et al. (2014). Nonstochastic reprogramming from a privileged somatic cell state. Cell 156, 649–662.

Henikoff, S., and Smith, M.M. (2015). Histone variants and epigenetics. Cold Spring Harb Perspect Biol 7, a019364.

Hossain, M., and Stillman, B. (2016). Opposing roles for DNA replication initiator proteins ORC1 and CDC6 in control of Cyclin E gene transcription. Elife 5.

Hu, X., Eastman, A.E., and Guo, S. (2019). Cell cycle dynamics in the reprogramming of cellular identity. FEBS Lett 593, 2840–2852.

Ivaldi, M.S., Karam, C.S., and Corces, V.G. (2007). Phosphorylation of histone H3 at Ser10 facilitates RNA polymerase II release from promoter-proximal pausing in Drosophila. Genes Dev 21, 2818–2831.

Jerkovic, I., Szabo, Q., Bantignies, F., and Cavalli, G. (2020). Higher-Order Chromosomal Structures Mediate Genome Function. J Mol Biol 432, 676–681.

Jiang, C., and Pugh, B.F. (2009). Nucleosome positioning and gene regulation: advances through genomics. Nat Rev Genet 10, 161–172.

Kalitsis, P., Zhang, T., Marshall, K.M., Nielsen, C.F., and Hudson, D.F. (2017). Condensin, master organizer of the genome. Chromosome Res 25, 61–76.

Kiger, A.A., Jones, D.L., Schulz, C., Rogers, M.B., and Fuller, M.T. (2001). Stem cell self-renewal specified by JAK-STAT activation in response to a support cell cue. Science 294, 2542–2545.

Kornberg, R.D. (1974). Chromatin structure: a repeating unit of histones and DNA. Science 184, 868–871.

Lai, W.K.M., and Pugh, B.F. (2017). Understanding nucleosome dynamics and their links to gene expression and DNA replication. Nat Rev Mol Cell Biol 18, 548–562.

Leatherman, J.L., and Dinardo, S. (2010). Germline self-renewal requires cyst stem cells and stat regulates niche adhesion in Drosophila testes. Nat Cell Biol 12, 806–811.

Li, V.C., Ballabeni, A., and Kirschner, M.W. (2012). Gap 1 phase length and mouse embryonic stem cell self-renewal. Proc Natl Acad Sci U S A 109, 12550–12555.

Li, V.C., and Kirschner, M.W. (2014). Molecular ties between the cell cycle and differentiation in embryonic stem cells. Proc Natl Acad Sci U S A 111, 9503–9508.

Lin, S., Yuan, Z.F., Han, Y., Marchione, D.M., and Garcia, B.A. (2016). Preferential Phosphorylation on Old Histones during Early Mitosis in Human Cells. J Biol Chem 291, 15342–15357.

Lipford, J.R., and Bell, S.P. (2001). Nucleosomes positioned by ORC facilitate the initiation of DNA replication. Mol Cell 7, 21–30.

Luger, K., Mader, A.W., Richmond, R.K., Sargent, D.F., and Richmond, T.J. (1997). Crystal structure of the nucleosome core particle at 2.8 A resolution. Nature 389, 251–260.

McCune, H.J., Danielson, L.S., Alvino, G.M., Collingwood, D., Delrow, J.J., Fangman, W.L., Brewer, B.J., and Raghuraman, M.K. (2008). The temporal program of chromosome replication: genomewide replication in clb5{Delta} Saccharomyces cerevisiae. Genetics 180, 1833–1847.

McGuffee, S.R., Smith, D.J., and Whitehouse, I. (2013). Quantitative, genome-wide analysis of eukaryotic replication initiation and termination. Mol Cell 50, 123–135.

Mendez, J., and Stillman, B. (2003). Perpetuating the double helix: molecular machines at eukaryotic DNA replication origins. Bioessays 25, 1158–1167.

Meshorer, E., and Misteli, T. (2006). Chromatin in pluripotent embryonic stem cells and differentiation. Nat Rev Mol Cell Biol 7, 540–546.

Monk, A.C., Siddall, N.A., Volk, T., Fraser, B., Quinn, L.M., McLaughlin, E.A., and Hime, G.R. (2010). HOW is required for stem cell maintenance in the Drosophila testis and for the onset of transit-amplifying divisions. Cell Stem Cell 6, 348–360.

Muller, C.A., Hawkins, M., Retkute, R., Malla, S., Wilson, R., Blythe, M.J., Nakato, R., Komata, M., Shirahige, K., de Moura, A.P., et al. (2014). The dynamics of genome replication using deep sequencing. Nucleic Acids Res 42, e3.

Nowak, S.J., and Corces, V.G. (2004). Phosphorylation of histone H3: a balancing act between chromosome condensation and transcriptional activation. Trends Genet 20, 214–220.

Paolinelli, R., Mendoza-Maldonado, R., Cereseto, A., and Giacca, M. (2009). Acetylation by GCN5 regulates CDC6 phosphorylation in the S phase of the cell cycle. Nat Struct Mol Biol 16, 412–420.

Patel, P.K., Arcangioli, B., Baker, S.P., Bensimon, A., and Rhind, N. (2006). DNA replication origins fire stochastically in fission yeast. Mol Biol Cell 17, 308–316.

Radman-Livaja, M., and Rando, O.J. (2010). Nucleosome positioning: how is it established, and why does it matter? Dev Biol 339, 258–266.

Raghuraman, M.K., Brewer, B.J., and Fangman, W.L. (1997). Cell cycle-dependent establishment of a late replication program. Science 276, 806–809.

Ramachandran, S., and Henikoff, S. (2016). Transcriptional Regulators Compete with Nucleosomes Post-replication. Cell 165, 580–592.

Randell, J.C., Bowers, J.L., Rodriguez, H.K., and Bell, S.P. (2006). Sequential ATP hydrolysis by Cdc6 and ORC directs loading of the Mcm2-7 helicase. Mol Cell 21, 29–39.

Ranjan, R., and Chen, X. (2021). Super-Resolution Live Cell Imaging of Subcellular Structures. J Vis Exp.

Ranjan, R., Snedeker, J., and Chen, X. (2019). Asymmetric Centromeres Differentially Coordinate with Mitotic Machinery to Ensure Biased Sister Chromatid Segregation in Germline Stem Cells. Cell Stem Cell 25, 666–681 e665.

Rhind, N., and Gilbert, D.M. (2013). DNA replication timing. Cold Spring Harb Perspect Biol 5, a010132.

Richmond, T.J., and Davey, C.A. (2003). The structure of DNA in the nucleosome core. Nature 423, 145–150.

Rodriguez, J., Lee, L., Lynch, B., and Tsukiyama, T. (2017). Nucleosome occupancy as a novel chromatin parameter for replication origin functions. Genome Res 27, 269–277.

Sawicka, A., and Seiser, C. (2012). Histone H3 phosphorylation - a versatile chromatin modification for different occasions. Biochimie 94, 2193–2201.

Serra-Cardona, A., and Zhang, Z. (2018). Replication-Coupled Nucleosome Assembly in the Passage of Epigenetic Information and Cell Identity. Trends Biochem Sci 43, 136–148.

Sheng, X.R., and Matunis, E. (2011). Live imaging of the Drosophila spermatogonial stem cell niche reveals novel mechanisms regulating germline stem cell output. Development 138, 3367–3376.

Shi, Z., Lim, C., Tran, V., Cui, K., Zhao, K., and Chen, X. (2020). Single-cyst transcriptome analysis of Drosophila male germline stem cell lineage. Development 147.

Simpson, R.T. (1990). Nucleosome positioning can affect the function of a cis-acting DNA element in vivo. Nature 343, 387–389.

Sivaguru, M., Urban, M.A., Fried, G., Wesseln, C.J., Mander, L., and Punyasena, S.W. (2018). Comparative performance of airyscan and structured illumination superresolution microscopy in the study of the surface texture and 3D shape of pollen. Microsc Res Tech 81, 101–114.

Snedeker, J., Wooten, M., and Chen, X. (2017). The Inherent Asymmetry of DNA Replication. Annu Rev Cell Dev Biol 33, 291–318.

Soufi, A., and Dalton, S. (2016). Cycling through developmental decisions: how cell cycle dynamics control pluripotency, differentiation and reprogramming. Development 143, 4301–4311.

Speck, C., Chen, Z., Li, H., and Stillman, B. (2005). ATPase-dependent cooperative binding of ORC and Cdc6 to origin DNA. Nat Struct Mol Biol 12, 965–971.

Speck, C., and Stillman, B. (2007). Cdc6 ATPase activity regulates ORC x Cdc6 stability and the selection of specific DNA sequences as origins of DNA replication. J Biol Chem 282, 11705–11714.

Stancheva, I. (2011). Revisiting heterochromatin in embryonic stem cells. PLoS Genet 7, e1002093.

Stewart-Morgan, K.R., Petryk, N., and Groth, A. (2020). Chromatin replication and epigenetic cell memory. Nat Cell Biol 22, 361–371.

Stillman, B. (2018). Histone Modifications: Insights into Their Influence on Gene Expression. Cell 175, 6–9.

Struhl, K., and Segal, E. (2013). Determinants of nucleosome positioning. Nat Struct Mol Biol 20, 267–273.

Sunchu, B., and Cabernard, C. (2020). Principles and mechanisms of asymmetric cell division. Development 147.

Szenker, E., Ray-Gallet, D., and Almouzni, G. (2011). The double face of the histone variant H3.3. Cell Res 21, 421–434.

Tanaka, T., Knapp, D., and Nasmyth, K. (1997). Loading of an Mcm protein onto DNA replication origins is regulated by Cdc6p and CDKs. Cell 90, 649–660.

Tarayrah, L., Li, Y., Gan, Q., and Chen, X. (2015). Epigenetic regulator Lid maintains germline stem cells through regulating JAK-STAT signaling pathway activity. Biol Open 4, 1518–1527.

Tazuke, S.I., Schulz, C., Gilboa, L., Fogarty, M., Mahowald, A.P., Guichet, A., Ephrussi, A., Wood, C.G., Lehmann, R., and Fuller, M.T. (2002). A germline-specific gap junction protein required for survival of differentiating early germ cells. Development 129, 2529–2539.

Tulina, N., and Matunis, E. (2001). Control of stem cell self-renewal in Drosophila spermatogenesis by JAK-STAT signaling. Science 294, 2546–2549.

Venkei, Z.G., and Yamashita, Y.M. (2015). The centrosome orientation checkpoint is germline stem cell specific and operates prior to the spindle assembly checkpoint in Drosophila testis. Development 142, 62–69.

Venkei, Z.G., and Yamashita, Y.M. (2018). Emerging mechanisms of asymmetric stem cell division. J Cell Biol 217, 3785–3795.

Vidaurre, V., and Chen, X. (2021). Epigenetic regulation of drosophila germline stem cell maintenance and differentiation. Dev Biol 473, 105–118.

White-Cooper, H., Leroy, D., MacQueen, A., and Fuller, M.T. (2000). Transcription of meiotic cell cycle and terminal differentiation genes depends on a conserved chromatin associated protein, whose nuclear localisation is regulated. Development 127, 5463–5473.

Wooten, M., Li, Y., Snedeker, J., Nizami, Z.F., Gall, J.G., and Chen, X. (2020a). Superresolution imaging of chromatin fibers to visualize epigenetic information on replicative DNA. Nat Protoc 75, 1188–1208.

Wooten, M., Ranjan, R., and Chen, X. (2020b). Asymmetric Histone Inheritance in Asymmetrically Dividing Stem Cells. Trends Genet 36, 30–43.

Wooten, M., Snedeker, J., Nizami, Z.F., Yang, X., Ranjan, R., Urban, E., Kim, J.M., Gall, J., Xiao, J., and Chen, X. (2019). Asymmetric histone inheritance via strand-specific incorporation and biased replication fork movement. Nat Struct Mol Biol 26, 732–743.

Xie, J., Wooten, M., Tran, V., Chen, B.C., Pozmanter, C., Simbolon, C., Betzig, E., and Chen, X. (2015). Histone H3 Threonine Phosphorylation Regulates Asymmetric Histone Inheritance in the Drosophila Male Germline. Cell 163, 920–933.

Yabuki, N., Terashima, H., and Kitada, K. (2002). Mapping of early firing origins on a replication profile of budding yeast. Genes Cells 7, 781–789.

Yadav, T., Quivy, J.P., and Almouzni, G. (2018). Chromatin plasticity: A versatile landscape that underlies cell fate and identity. Science 361, 1332–1336.

Yadlapalli, S., Cheng, J., and Yamashita, Y.M. (2011). Drosophila male germline stem cells do not asymmetrically segregate chromosome strands. J Cell Sci 724, 933–939.

Yadlapalli, S., and Yamashita, Y.M. (2013). Chromosome-specific nonrandom sister chromatid segregation during stem-cell division. Nature 498, 251–254.

Yamashita, Y.M., Jones, D.L., and Fuller, M.T. (2003). Orientation of asymmetric stem cell division by the APC tumor suppressor and centrosome. Science 301, 1547–1550.

Yamashita, Y.M., Mahowald, A.P., Perlin, J.R., and Fuller, M.T. (2007). Asymmetric inheritance of mother versus daughter centrosome in stem cell division. Science 315, 518–521.

Yim, H., and Erikson, R.L. (2010). Cell division cycle 6, a mitotic substrate of polo-like kinase 1, regulates chromosomal segregation mediated by cyclin-dependent kinase 1 and separase. Proc Natl Acad Sci U S A 107, 19742–19747.

Yu, C., Gan, H., Han, J., Zhou, Z.X., Jia, S., Chabes, A., Farrugia, G., Ordog, T., and Zhang, Z. (2014). Strand-specific analysis shows protein binding at replication forks and PCNA unloading from lagging strands when forks stall. Mol Cell 56, 551–563.

Yuan, Z., Schneider, S., Dodd, T., Riera, A., Bai, L., Yan, C., Magdalou, I., Ivanov, I., Stillman, B., Li, H., et al. (2020). Structural mechanism of helicase loading onto replication origin DNA by ORC-Cdc6. Proc Natl Acad Sci U S A 117, 17747–17756.

Zhang, W., Feng, J., and Li, Q. (2020). The replisome guides nucleosome assembly during DNA replication. Cell Biosci 10, 37.

Zhang, Y., Sun, Z., Jia, J., Du, T., Zhang, N., Tang, Y., Fang, Y., and Fang, D. (2021). Overview of Histone Modification. Adv Exp Med Biol 1283, 1–16.

Zion, E.H., Chandrasekhara, C., and Chen, X. (2020). Asymmetric inheritance of epigenetic states in asymmetrically dividing stem cells. Curr Opin Cell Biol 67, 27–36.

