## Supplemental Information for "Differential condensation of sister chromatids coordinates with Cdc6 to ensure distinct cell cycle progression in *Drosophila* male germline stem cell lineage"

**Materials and Methods**

### 1. KEY RESOURCES TABLE

| REAGENT or RESOURCE | SOURCE | IDENTIFIER |
|---|---|---|
| **Antibodies** | | |
| Mouse monoclonal anti-Fascilin III (Fas III) | Developemental Studies Hybridoma Bank (DSHB) | RRID: AB_528238 |
| Mouse monoclonal anti-alpha Spectrin | Developemental Studies Hybridoma Bank (DSHB) | RRID: AB_528473 |
| Mouse monoclonal anti-Armadillo | Developemental Studies Hybridoma Bank (DSHB) | RRID: AB_528089 |
| Rat monoclonal anti-DE-cadherin | Developemental Studies Hybridoma Bank (DSHB) | RRID: AB_528120 |
| Mouse monoclonal anti-lamin | Developemental Studies Hybridoma Bank (DSHB) | RRID: AB_528336 |
| Rabbit monoclonal anti-H3T3ph | Millipore | 05-746R |
| Rabbit anti-Vasa | Santa Cruz | SC-30210 |
| Mouse monoclonal anti-H3S10ph | Abcam | AB-14995 |
| Guinea pig anti-Traffic jam | Kindly provided by Mark Van Doren, Johns Hopkins University, MD, USA | N/A |
| Mouse anti-PCNA | Santa Cruz | Santa Cruz sc-56 |
| Chicken anti-GFP | Abcam | Abcam ab 13970 |
| **Experimental Models: Transgenic Organisms/Strains** | | |
| *D. melanogaster*. UAS-α-tubulin-GFP | Bloomington *Drosophila* Stock Center | RRID: BDSC_7373 |

| | | |
|---|---|---|
| *D. melanogaster*. hs-flp | Bloomington *Drosophila* Stock Center | RRID: BDSC_26902 |
| *D. melanogaster*. Nos-Gal4 | Kindly provided by Mark Van Doren, Johns Hopkins University, MD, USA | N/A |
| *D. melanogaster*. Nos-Gal4; tubulin-Gal80$^{ts}$ | Kindly provided by Yukiko Yamashita, Whitehead Institute, MIT, USA | N/A |
| *D. melanogaster*. UAS-H3-mCherry-H3-EGFP or UAS-H3- EGFP-H3-mCherry | Our lab has developed | N/A |
| *D. melanogaster*. UAS-H4-mCherry-H4-EGFP | Our lab has developed | N/A |
| *D. melanogaster*. UAS-H2A-mCherry-H2A-EGFP | Our lab has developed | N/A |
| UASp-YFP-PCNA | Kindly provided by Patrick O’Farrell, UCSF, San Francisco, | N/A |
| pcna>PCNA-eGFP | Kindly provided by Shelby Blythe and Eric Wieschaus | N/A |
| **Experimental Models: Knock-in Organisms/Strains** | | |
| *D. melanogaster*. Cdc6-mCherry | Our lab has developed | N/A |
| **Chemicals** | | |
| FBS | Sigma | Cat#F2442 |
| Penicillin/Streptomycin | Sigma | Cat#P0781 |
| Nocodazole | Sigma | Cat#M1404 |
| Colcemid | Sigma | 10295892001 Roche |
| DMSO | Sigma | Cat#D2650 |

| | | |
|---|---|---|
| Mounting Medium | Vector | Cat# H-1400 |
| BSA | Cell Signaling Technology | Cat#9998 |
| Formaldehyde | Fisher Scientific | Cat#F79-500 |
| Schneider Medium | ThermoFisher Scientific | Cat# 21720024 |
| Insulin | ThermoFisher Scientific | Cat#12585014 |
| FluoroDish | World Precision Instrument, Inc. | Cat#FD35PDL |
| **Software and algorithms** | | |
| Adobe Illustrator CS6 | Adobe | N/A |
| Prism 6 | Graphpad | N/A |
| Fiji | NIH | N/A |
| EndNote | Clarivate Analytics | N/A |
| IMARIS | Bitplane | N/A |

### 2. FLY STRAINS AND HUSBANDRY

#### Transgenic fly strains

Fly stocks were raised using standard Bloomington medium at 18°C, 25°C, or 29°C as noted. The following fly stocks were used: *hs-flp* on the X chromosome (Bloomington Stock Center BL-26902), *nos-Gal4* on the 2nd chromosome (Van Doren et al., 1998), *UASp-FRT-H3-GFP-PolyA-FRT-H3-mKO* on the 3$^{rd}$ chromosome as reported previously (Tran et al., 2012), *UASp-FRT-H3-mCherry -PolyA-FRT-H3-eGFP* on the 3$^{rd}$ chromosome, *UASp-FRT-H4-mCherry -PolyA-FRT-H4-eGFP* on the 3$^{rd}$ chromosome, *UASp-FRT-H2A-mCherry -PolyA-FRT-H2A-eGFP* on the 3$^{rd}$ chromosome, *UAS-$\alpha$-Tubulin-GFP (*Bloomington Stock Center BL-7373*)* on the 3$^{rd}$ chromosome, *UASp-FRT-H3S10A-mCherry -PolyA-FRT-H3S10A-eGFP* on the 3$^{rd}$ chromosome, w; UASp-YFP-PCNA/CyO (McCleland et al., 2009) (from Patrick O'Farrell, Department of Biochemistry and Biophysics, University of California, San Francisco, USA) and *nos-Gal4 (without VP16)/Cyo; tub-Gal80$^{ts}$/TM6B* (from Yukiko Yamashita, University of Michigan, Ann Arbor, Michigan, USA).

**Generating knock-in fly strains**

In collaboration with Fungene Inc. (Beijing, China), the following fly lines were generated using the CRISPR-Cas9 technology: CG5971 (*cdc6*) with mCherry tag at the N-terminus, in order to generate the following fusion protein: CDC6-mCherry N term.

### 3. METHOD DETAILS

**Immunostaining**

Immunofluorescence staining was performed using standard procedures (Hime et al., 1996; Tran et al., 2012). For immunofluorescence staining, testes from young (0-2 day old) flies were dissected in PBS buffer. Samples were then fixed in 4% formaldehyde in phosphate-buffered saline (PBS) for 10 minutes at room temperature, washed thrice for 15 minutes per wash in PBST (PBS with 0.1% Triton X-100) and blocked for at least 30 minutes in PBST with 3% bovine serum albumin at room temperature. Samples were incubated for overnight (~16 hours) at 4°C in primary antibodies. Then sample were washed thrice for 15 minutes per wash in PBST and incubated in a 1:1000 dilution of Alexa Fluor-conjugated secondary antibody (from Molecular Probes) in PBST for 2 hours at room temperature. Sample were washed quickly three times with PBST and then four times 10 minutes each with PBST. Samples were mounted for microscopy in Vectashield antifade mounting medium (Vector Laboratories, Cat# H-1400) with/without DAPI, and examined using Leica SPE confocal microscope, Zeiss LSM 700 confocal microscope, or Zeiss LSM 800 confocal microscope with Airyscan with 63x oil immersion objectives. Images of separate fluorochromes from multiple stained tissues were collected individually and combined using Fiji and Adobe illustrator (Adobe).

**Heat shock scheme**

Flies with *UASp*-dual color histone transgenes were paired with the *nos-Gal4* driver. Flies were raised at 18°C throughout development until adulthood to avoid pre-flip (Tran et al., 2012). Before heat shock, 1-to-3-day old males were transferred to vials that had been air dried for 24 hours. Vials were submerged underneath water up to the plug in a circulating 37°C water bath for 90 minutes and recovered in a 29°C incubator for indicated time before dissection for immunostaining experiments.

**PCNA-YFP detection**

Male flies expressing containing the genotype UASp-YFP-PCNA/CyO (McCleland et al., 2009) were crossed with virgin females expressing the *nos-Gal4* driver. Crosses were maintained at 18°C during development. Upon eclosion, progeny were transferred to room temperature for 1-2 days. Testes were then dissected in Schneider's Drosophila medium (Gibco, catalog # 21720001) at room temperature (RT) and incubated in Schneider's medium containing 10 μM EdU analog (Invitrogen Click-iT EdU Imaging Kit, catalog # C10340). Testes were incubated for 10 minutes, rotating at RT. At the end of the 10 minutes, testes were washed three times with Schneider's medium at RT. Testes were then fixed, immunostained and mounted as described above.

**Chromatin fiber assay and quantification**

For a full description of the protocol please refer to (Wooten et al., 2020). In brief, testes were dissected in Schneider's *Drosophila* media (Gibco, Catalog # 21720001) then incubated in 10 μM EdU for 15 minutes at room temperature. The Schneider's media was then washed off and testes were transferred to lysis buffer (100 mM NaCl, 25 mM Tris-base, 0.2% Joy detergent, pH 10). The testes were then transferred by pipette to a slide with excess lysis buffer. Excess lysis buffer was then removed by Kimwipe and the early germ cells were isolated by microdissection of the end of the testis tip. This works by making an incision at the end of the testis membrane and then allowing the early germ cells to spill out of the tissue before manually removing the rest of the tissue. At the end of the microdissections (~5 min), the lysis buffer solution should be nearly dry. At this point 20 μL of lysis buffer is added back to the lysing cells. After 5 minutes, 10 μL of a sucrose/formalin (1M sucrose; 10% formaldehyde) mixture is added to the lysis buffer solution and allowed to sit for another minute. At this point, a 24x60 coverslip (Fisher brand Microscope Cover Glass, 12-545-J 24 × 60 mm) is very slowly set down on top of the solution to spread fibers. The slide is the moved to -80 °C for 45 min to freeze. After fully freezing, the coverslip is quickly removed by razor blade and the slide is transferred to 95% EtOH for 10 minutes at -20 °C. The slides are then incubated for 1 min in fixative solution (0.5% formaldehyde in 1× PBST (1× PBS with 0.1% Triton)). After draining the fixative solution, slides are then washed three times in Coplin jars with PBST. The slides were then quickly dried and transferred to a humidity chamber and primary antibodies were then added overnight and the slides are covered with parafilm. After washing three times, the slides were then transferred back to the humidity chamber and incubated for two hours with secondary antibodies. To label Edu, click chemistry is then performed to adhere a 637-Biotin-Azide dye to the previously incorporated EdU using a kit (Life Science C10640).

For the total histone asymmetry work, unflipped histone labeled strains are generated by crossing a FRT-Histone-GFP-FRT-Histone-mCherry line with nanos-Gal4 without hs-flp. The chromatin fibers are prepared from these lines as described. For these fibers, the primary antibodies are anti-PCNA (1:200, Santa Cruz sc-56) and anti-GFP (1:1,000; Abcam ab 13970). The slides are mounted with ProLong Diamond mounting media with Dapi. The fibers were imaged using Airyscan super-resolution imaging. The images are analyzed on FIJI ImageJ by measuring 2μm segments of replicating DNA on either sister chromatid in a replicating region and the ratio of these values is calculated as $Log_2$ ((Leading Strand Average Intensity – Background Average Intensity)/ (Lagging Strand Average Intensity – Background Average Intensity)). The lagging strand is identified by the preferential incorporation of PCNA on that side.

### 4. MICROSCOPY

#### Live cell imaging

To examine the temporal dynamics of cellular processes during asymmetric GSC divisions, we conducted live cell imaging with high temporal resolution (e.g. 1min or 2min interval as mention in the supplemental movie legend). To perform live cell imaging, adult *Drosophila* testes were dissected in a medium containing Schneider's insect medium with 200 μg/ml insulin, 15% (vol/vol) FBS, 0.6x pen/strep, with pH value at approximately 7.0, which we called "live cell medium" as reported previously (Ranjan et al., 2019). Testes were then placed on a Poly-D-lysine coated FluoroDish (World Precision Instrument, Inc.), which contains the live cell medium as described. All movies were taken using spinning disc confocal microscope (Zeiss) equipped with an evolve-TM camera (Photometrics), using a 63x Zeiss objective (1.4 NA) at

29°C. The ZEN 2 software (Zeiss) was used for acquisition with 2x2 binning. All videos for live cells are shown in Movie S1 to S8.

**Super-Resolution Live Snapshot (SRLS)**

To achieve high spatial and temporal resolution to understand dynamics of cellular processes during asymmetric GSC divisions, we conducted Super-Resolution Live Snapshot (SRLS). For a full description of the protocol please see [(Ranjan and Chen, 2021), Figure S3B]. In brief, adult *Drosophila* testes, expressing CDC6-mCherry were dissected in live cell medium (containing Schneider's insect medium with 200 µg/ml insulin, 15% (vol/vol) FBS, 0.6x pen/strep, with pH value at approximately 7.0). Testes were then placed on a Poly-D-lysine coated glass bottom FluoroDish (World Precision Instrument, Inc.), which contains the live cell medium as described. Testes were incubated at room temperature for ~ 20-30 min before imaging to stabilize the tissue in ex-vivo condition and to minimize the interference due to movement. All SRLS was taken using Zeiss LSM 800 confocal microscope with AiryScan super-resolution module equipped with highly sensitive GaAsP (Gallium Arsenide Phosphide) detectors using a 63x Zeiss objective (1.4 NA) at ~20°C.

**Fixed cell imaging**

To examine the spatial dynamics of cellular processes and spatial localization of cellular component during asymmetric GSC divisions, we conducted fix cell imaging with high spatial resolution. After immunostaining, images were taken using Zeiss LSM 700 confocal microscope, or Zeiss LSM 800 confocal microscope with Airyscan super-resolution mode with 63x oil immersion objectives. To avoid interference of free-floating histones in the quantification, fixed cell imaging was performed to investigate the differential condensation of old and new histone enriched chromatin in control and loss of differential condensation in H3S10A mutant. Images were processed using Imaris software (3D image reconstruction) and Fiji software (to quantify the total amount of protein "RawIntDen" or to generate maximum intensity projection).

### 5. QUANTIFICATION AND STATISTICAL ANALYSIS

#### The 3D quantification of movies and fix images

To quantify total amount of protein (such as histones or CDC6) during asymmetric GSC divisions and symmetric SG divisions, we conducted a 3D quantification (quantification in volume) by measuring the fluorescence signal in each plane from the Z-stack (e.g. Figure S1B) as earlier (Ranjan et al., 2019). The 3D quantification was done at different cell cycle stages (as labeled in the corresponding Figures) in GSCs and SGs (e.g. 8-cell cyst), using time lapse movie with *cdc6-mCherry* knock-in lines *and H3-mCherry, H4-mCherry, H2A-mCherry,* transgenic lines. No antibody was added to enhance EGFP or mCherry signals for quantification.

The fixed immunostaining images used fluorescence signal of mCherry or EGFP (histone, PCNA or CDC6), or Edu (thymidine analogue). The live cell images used fluorescence signal of mCherry or EGFP (histone, PCNA or CDC6).

The 3D quantification of the fluorescence signal was done manually using Fiji (Image J). Un-deconvolved raw images as 2D Z-stacks were saved as un-scaled 16-bit TIF images, and the sum of the gray values of pixels in the image ("RawIntDen") was determined using Fiji (Image J). A circle was drawn to include all fluorescence signal (marked by EGFP or mCherry), and an identical circle was drawn in the hub region as the background. The gray values of the fluorescence signal pixels for each Z-stack (Foreground signal, Fs) was calculated by subtracting the gray values of the background signal pixels (Background signal, Bs) from the gray values of the raw signal pixels (Raw signal, Rs). The total amount of the fluorescence signal in the nuclei was calculated by adding the gray values of the fluorescence signal from all Z-stacks. The total amount of the fluorescence signal (Fs) in the nuclei with Z-stacks (Z1+…+Zn): Fs (Z1+…+Zn) = [(Rs-Bs)1+…+(Rs-Bs) n]. The total amount of fluorescence signal or gray value would represent the total amount of protein in the cell (quantification in volume of the cell). This method is also referred to in text as the sum of slices method.

**Nucleosome density assay**

For nucleosome density quantification, first we calculated the total amount of histone in the nucleus [quantification in volume (3D quantification)], this would represent the amount of histones that presents on entire length of DNA at that cell cycle stage. Two copy of each core histones (2x H3, 2x H4, 2x H2A and 2x H2B) are present in the nucleosome. Therefore, knowing total amount of histone would allow us to calculate global nucleosome density on DNA; the calculated nucleosome density = ½ amount of histone/DNA length (Figure S1B). Then, we determined the ratio of nucleosome density between daughter cells in telophase (GSC/GB or SG1/SG2) to understand nucleosome density pattern of sister chromatids during asymmetric stem cell division. For global nucleosome density quantification, we have used histone H3, H4 and H2A. We have looked at telophase stage where sister chromatids are separated and individual sister chromatids can be visualized. Further, in telophase stage new histone incorporation would not occur, therefore nucleosome density could be analyze accurately.

We calculated the global nucleosome density ration of GSC side to GB side chromatids in telophase. The nucleosome density ratio $(GSC/GB)_{telophase}$ = ½ total amount of histone in GSC side/DNA length / ½ total amount of histone in GB side/DNA length = total amount of histone in GSC side / total amount of histone in GB side. This gives a relative difference in nucleosome density in the daughter cells in telophase. We observed that a ~1.50-fold high H3 nucleosome density on sister chromatids in the GSC side compared to the GB side.

We have used similar approach to determine nucleosome density of the sister chromatids immediately after DNA replication using chromatin fiber technique to understand whether asymmetry in nucleosome density has been established during DNA replication. Indeed, consistent with the global nucleosome density asymmetry in telophase, we observed that the leading strand has higher histone H3 nucleosome density compared to the lagging strand. Further, quantification showed a ~1.44-fold H3 nucleosome density asymmetry after DNA replication which is comparable with the global nucleosome density asymmetry in telophase.

**A quantitative assay for sister chromatids condensation**

To quantitatively monitor sister chromatids condensation, we used an area-based method as previously described (Maddox et al., 2006). A dual-color histone labeled (UASp-FRT-histone-mCherry-PolyA-FRT-histone-EGFP-PolyA) *Drosophila* melanogaster transgenic lines were used for this purpose, as described earlier [(Tran et al., 2012), Figure S1A]. The GSC or the SG with old and new histones were imaged, and all condensation assay was performed using FIJI (ImageJ) software. A maximum intensity projection was generated for each GSCs or SGs. The largest square region that fit within the GSCs or SGs nucleus was cut out for analysis (Figure S2H). Next, the intensity of each pixel of the image was determined using "display_pixel_values" plugins using FIJI (ImageJ) software. Then, the images of the GSC or SG nucleus were individually scaled, setting the minimum intensity to 0 and the maximum to 65,535

(16-bit range). Individual scaling of the image ensures that the fluorescence intensity distribution is independent of fluctuations due to changes in illumination intensity, variations between the sample, or photobleaching.

Sister chromatids condensation is accompanied by a progressive change in the shape of the fluorescence intensity distribution of the nuclear histone-EGFP and histone-mCherry signal (Fig). Prior to the mitotic onset and chromatin condensation (such as in interphase), the old histone-EGFP and new histone-mCherry fluorescence signal are relatively homogeneous. However, as the sister chromatids condense, the old histone-EGFP and new histone-mCherry fluorescence concentrate in a smaller area of the image. To quantify this shift in the fluorescence intensity distribution and the difference in old and new histone enriched chromatin condensation dynamics, we monitored the pixels across the image with a threshold at 35% of the maximum intensity of the image (scaled intensities: 22,937). Condensation kinetic profiles to compare old histone enriched chromatids and new histone enriched chromatids were generated by plotting the percentage of pixels below the threshold (the condensation parameter). For every condition (control or perturbation), the condensation parameters measured of the prometaphase GSCs or SGs were averaged and plotted. Averaging minimizes the contribution of spatial fluctuations. Quantification shows, in GSCs, old and new histone enriched sister chromatids show a differential condensation, whereas, in the symmetrically dividing spermatogonia cells, both sister chromatids condense symmetrically.

**Quantification of mitotic Cdc6 chromatin association**

To enrich for prometaphase and metaphase cells, flies were treated with 4 μg/mL colcemid for four hours after dissection in imaging media with insulin and left at RT in the dark. Afterwards, standard Immunofluorescent labeling was performed. In FIJI, using a single slice of the cell encapsulating the largest volume of chromatin, outline the chromatin occupied region using DAPI signal (Figure S3A). We then measured the average intensity and area of Cdc6 in this region. We repeated this process to measure the area and Cdc6 intensity throughout the entire slice of the cell. The relative amount of Cdc6 in the chromatin occupied region is calculated by multiplying the area of the chromatin occupied region by the average intensity of Cdc6 present in the region. The relative amount of Cdc6 excluded from the chromatin region is calculated by multiplying the area of the entire cell by the average intensity of Cdc6 throughout the entire cell and then subtracting out the relative amount of Cdc6 that is in the chromatin occupied region. To get a ratio of these amount and therefore the relative portion of Cdc6 that is chromatin bound, simply divide the relative chromatin occupied amount by the relative chromatin excluded amount. This analysis is done using a single slice as it can be hard to define the upper and lower bounds of the cell membrane. This does mean that the ratio is simply a fraction of the signal in that slice associated with chromatin rather than the true fraction of protein associated with DNA. However, this is still a very useful approach for defining the changes in protein association with mitotic chromatin over time.

**Analysis of post-mitotic early replicating pairs**

All imaging analysis and quantification was performed using the imaging processing software FIJI. Post-mitotic GSC-GB pairs were identified by their stereotypical perpendicular division plane from the hub, their temporarily shared cytoplasm and/or the presence of the spectrasome between cells. Timing of replication was approximated using nucleus size. The nucleus of a cell is maximally compacted at telophase and gradually decompacts in G1/S. Nuclear size continues to increase throughout S-phase as DNA content increases. Therefore, the earliest replicating cells are markedly smaller than cells in mid-S phase and late-S phase. As such, nucleus size can be used as a measure of time after division. For single slice measurements (Figure 4), we assessed nucleus size by calculating the maximum area occupied by a nucleus in a single slice of a z-stack image. For each slice of the z-stack image, area was assessed using FIJI by drawing a circle around the edge of the nucleus. Once the largest slice was identified, the region contained within the circle was then measured for fluorescence levels of PCNA-YFP and EdU.  For sum of slices measurements in Figures 5-6, nucleus size is approximated by summing the area from all total slices and corrected by multiplying by the Z-stack size. Total fluorescence in this context is similarly calculated by summing total fluorescence from each slice. For the sum of slice approach used with the Nocodazole (NZ) disruption, PCNA-EGFP was used (Blythe and Wieschaus, 2016). EdU was measured for the H3S10A disruption assay to be compatible with labeling old and new histones. All values are reported as the ratio of the intensity of PCNA or EdU in the GSC over the GB. Thus, negative values in the earliest cells represent the GB entering replication prior to the GSC.

**Mitotic index**

Mitotic index was analyzed by using a mitotic specific epigenetic mark, histone H3 threonine 3 phosphorylation (H3T3ph). The number of H3T3ph positive GSCs and total GSCs were counted in control testes and testes with perturbations. The percentage of H3T3ph positive GSCs were calculated that reflect the mitotic index of the testes. Mitotic index = Number of H3T3ph positive GSCs / Total GSCs x 100.

**Analysis of centrosome mis-orientation**

GSCs expressing either H3 or H3S10A are scored. Centrosomes are labeled using anti-γ-Tubulin. Centrosomes are oriented with the mother centrosome located near the hub-GSC interface, while the daughter centrosome moves toward the opposite side during G2 phase (Yamashita et al., 2003; Yamashita et al., 2007). Mother centrosome is anchored by astral microtubule toward the hub-GSC interface (Yamashita et al., 2007), therefore in normal GSCs one centrosome should always be adjacent to the GSC-hub interface. In GSCs where both centrosomes are positioned away from the GSC-hub interface, it is defined as misoriented centrosomes (Yamashita et al., 2007).

**Co-localization assay**

The co-localization assay was performed using FIJI (ImageJ) software. The image was imported into ImageJ, and the GSCs or SGs that are in late prophase or prometaphase were selected. We draw a box around GSC or SG of interest using the "box tool" on the toolbar. We use chromatin-bound histone signals to draw the box, and the box should be as tight to the edges of the cell as possible. After drawing the box, the image was duplicated using an option available under the "image" tab on the toolbar (Image>Duplicate Image). Next, channels were split using an option available under the "image" tab on the toolbar (select Image>Color>Split Channels). Now all channels are available to perform all co-localization permutations possible. To perform co-localization analysis of H3S10ph with old or new histone enriched chromatin, Coloc 2 plugin was used (select Analysis>Co-localization>Coloc 2). The Coloc 2 implements and performs the pixel intensity correlation over space (pixel intensity spatial correlation analysis). The analysis was performed on relevant channel combinations such as S10P vs. old Histone and S10P vs. new

Histone. This generates a pdf file, which summarizes the results of the co-localization analysis. We use the Spearman value to compare co-localization between different datasets. The result is +1 for perfect correlation, 0 for no correlation, and -1 for perfect anti-correlation.

**Defining different categories of asymmetry**

To define different categories (asymmetric and symmetric) for nucleosome density, histone segregation, and CDC6 segregation, we used those ratios in symmetrically dividing SGs (SG1/SG2) to define the symmetric range. For example, H2A nucleosome density in telophase, ratios above mean + SE (1.059 + 0.036 = 1.095) are called ‘asymmetric’ and below are called ‘symmetric’.

**Statistics**

Statistics analysis was performed in Prism 6 (GraphPad) and was done with Mann-Whitney unpaired t-test and one sample t-tests. Data are presented as Average ± SE and significant difference between two groups were noted by asterisks (**** $p < 0.0001$).

### 6. DISRUPTION ASSAYS

**Compromising H3 Serine 10 Phosphorylation by introducing non-phosphorylable Alanine (serine to alanine)**

The *UAS-H3-mCherry* and *UAS-H3S10A-mCherry* flies were crossed with *nanos-Gal4* (*without VP16*)*; tub-Gal80ts* flies to generate *tub-Gal80ts, nos-Gal4>H3-mCherry* (*Ctrl*), and *tub-Gal80ts, nos-Gal4> H3S10A-mCherry* (H3S10A mutant) male flies. Crosses were maintained at 18°C (permissive temperature for Gal80ts) to keep the Gal4 repressed; therefore, no H3S10A expression during development. After eclosion, flies were shifted to 29°C (restrictive temperature for Gal80ts) to allow the Gal4 expressed, therefore, H3S10A expression for 0-5 days (as indicated).

**Disruption of microtubule asymmetry**

Temporal asymmetry in microtubes activity was disrupted using previously standardized conditions in the *Drosophila* male GSCs (Ranjan et al., 2019). To disrupt the temporal asymmetry of microtubules activity at mother and daughter centrosomes, testes were briefly (2-3 hours) treated with microtubule depolymerizing drug nocodazole (NZ), and then NZ was washed out to allow cells to resume the cell cycle. Previously we showed that #1. This disrupts pre-established microtubule asymmetry in GSCs, and #2. When cell resume cell cycle after NZ wash, microtubules activity at mother and daughter centrosome becomes nearly symmetric (Ranjan et al., 2019). Here, we considered that when we allow the cell to progress into the next cell cycle after NZ wash, asymmetric microtubules activity at mother and daughter centrosome will re-established.

Prior to dissection, Nocodazole (NZ) solution was freshly prepared by adding 1 µl of 2 mg/ml stock solution of NZ in DMSO per 200 µl of "live cell media" for a final NZ concentration at 10µg/ml. This solution was left in the dark at room temperature (RT) until needed. Testes were dissected in live-cell media and transferred to tubes with the excess live-cell media. After removing the live-cell media, 50 µl of NZ solution was added to each tube, which was left open in darkness at RT for 2-4 hours in the NZ solution. At the end of the 2-4 hours, the NZ solution was removed, and tissue was wash in live-cell media. The media was removed, and fresh media was added, repetitively for a total of at least three washes within 5 minutes and incubate the tissue in the live cell media as required (Figure 5A). For the PCNA and EdU experiments, testes were then fixed approximately 50-60 minutes after release to catch anaphase or telophase cells. For the CDC6 SRLS and H3 segregation experiments, testes were live imaged immediately after NZ release (for H3 segregation experiments and cdc6 experiments), and for the H3 recovery experiment, testes were live imaged 4-5 hours after NZ release.

**Supplementary figures and figure legends:**

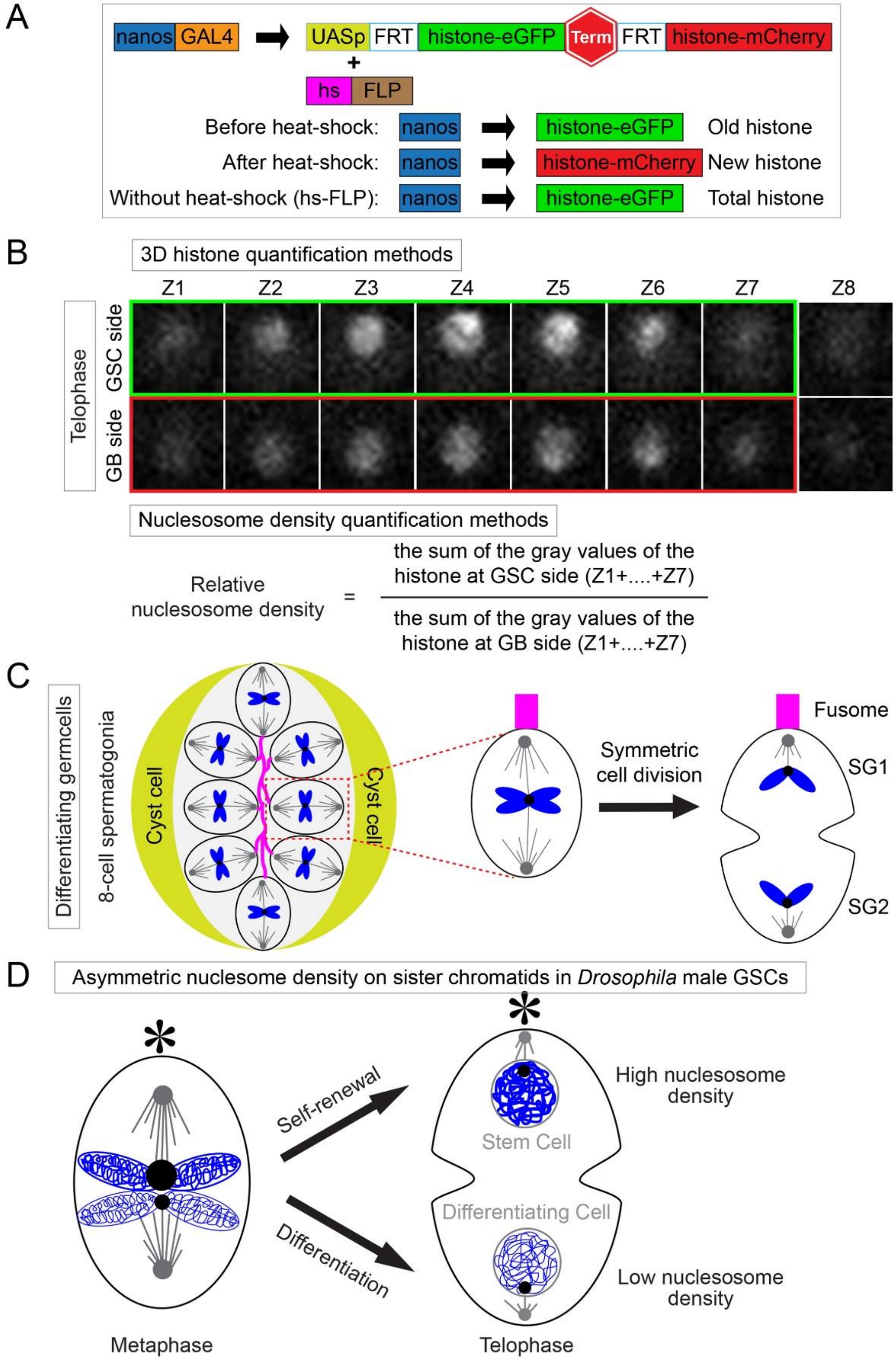

**Figure S1.** (**A**) Cartoon depiction of the tag switching approach for differential labeling of old and new histones, and total histone, as well as the *UASp-FRT-histone-eGFP-PolyA-FRT-histone-mCherry-PolyA* transgene, similar to (Tran et al., 2012). UAS, upstream activating sequence; FRT, FLP (flippase) recombination target; histone, H3, H4 and H2B driven by *nanos-Gal4*, a germline-specific driver; hs-FLP, the yeast FLP recombinase controlled by the heat shock (hs) promoter. (**B**) Description of the sum of slices quantification method for calculating relative nucleosome density after division. (**C**) Description of the method used to define the two SGs for quantification purposes based on proximity to the fusome. (**D**) Model describing the nucleosome density asymmetry generated during GSC ACD.

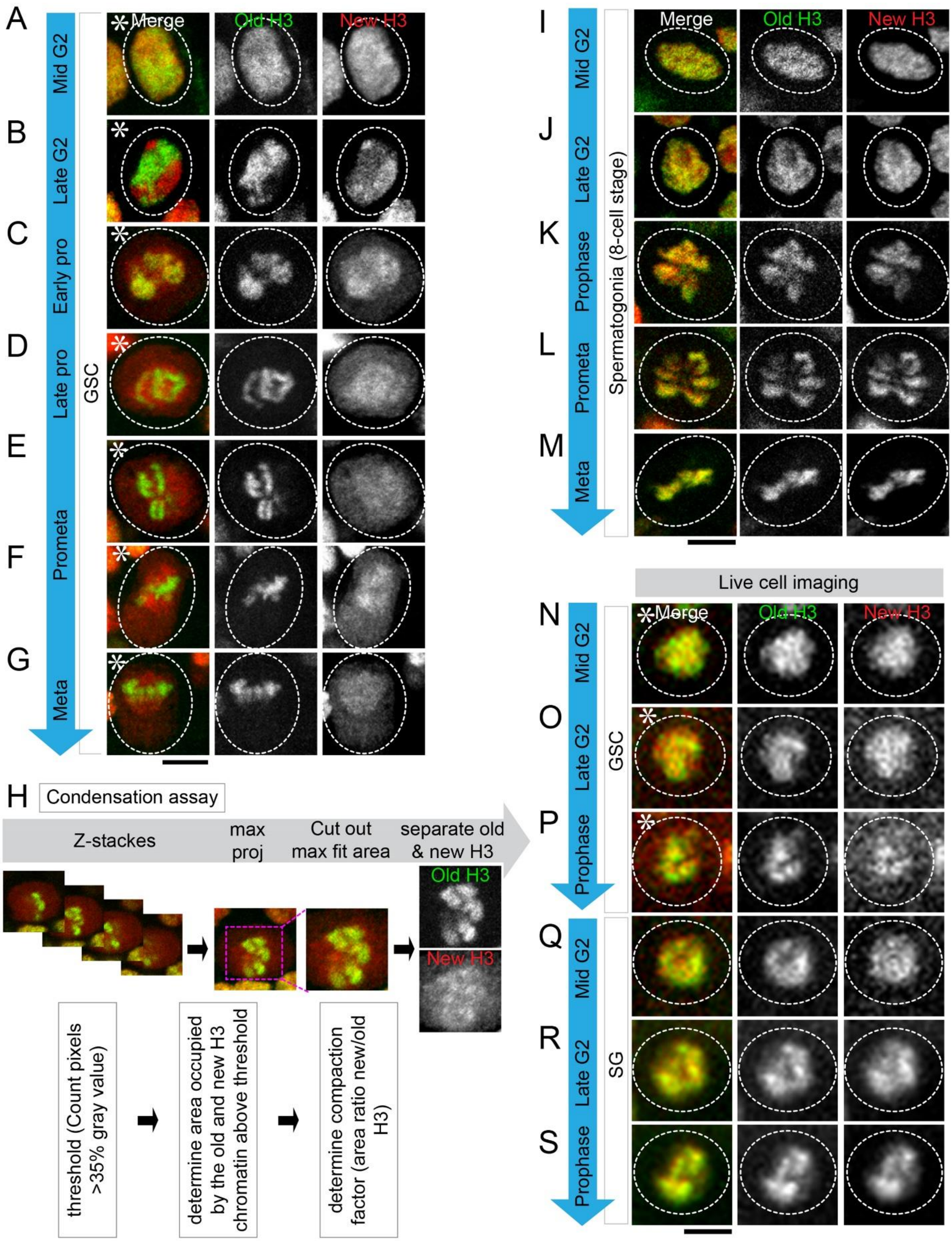

A
B
C
D
E
F
G
Mid G2
Late G2
Early pro
Late pro
Prometa
Meta
GSC
Merge
Old H3
New H3
I
J
K
L
M
Mid G2
Late G2
Prophase
Prometa
Meta
Spermatogonia (8-cell stage)
H
Condensation assay
Z-stackes
max proj
Cut out max fit area
separate old & new H3
threshold (Count pixels >35% gray value)
determine area occupied by the old and new H3 chromatin above threshold
determine compaction factor (area ratio new/old H3)
Live cell imaging
N
O
P
Q
R
S
Prophase
SG

**Figure S2.** (**A-G**) A detailed high-resolution progression of differential condensation throughout mitosis showing that old histone enriched chromatids condense first and remained more condensed through metaphase in GSCs. (**H**) Description of the method used for quantifying the compaction factor of old and new H3 by thresholding. (**I-M**) A detailed high-resolution progression of symmetric condensation throughout mitosis showing that old and new histone enriched chromatids condense equally through metaphase in SGs. (**N-S**) Live Cell imaging of the early stages of compaction showing that the separation of old and new histones becomes distinguishable in late G2 and differential condensation of old and new histone enriched chromatids in prophase in GSCs (**N-P**), but not in SGs (**Q-S**). Scale bars are 2 μm. Asterisk: hub.

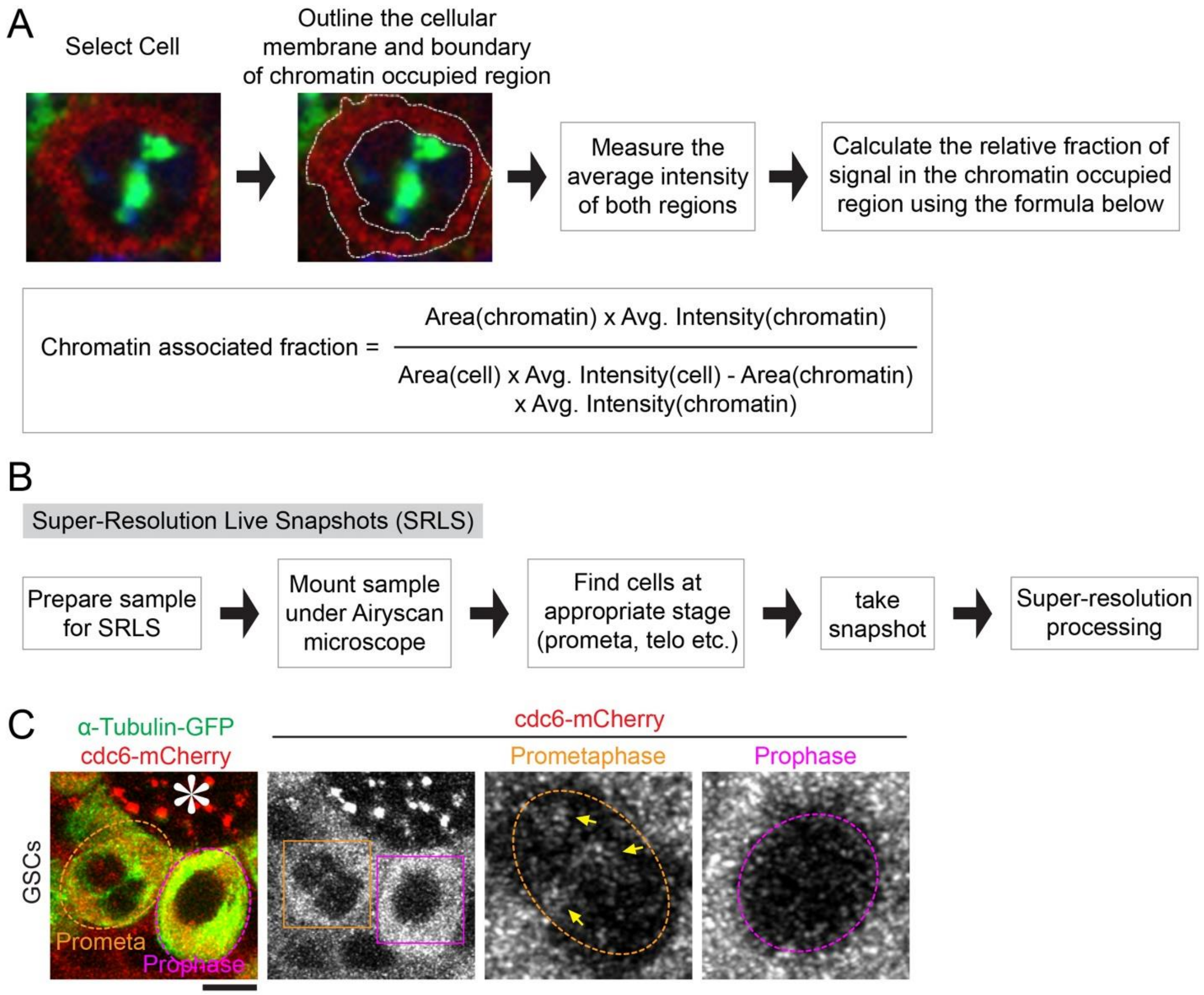

**Figure S3.** (**A**) Description of the method used to quantify the relative chromatin association of Cdc6 during mitosis. (**B**) Description of the protocol used for Super-Resolution Live Snapshots (SRLS). (**C**) Image of prophase and prometaphase GSCs using SRLS. Scale bars are 5 μm. Asterisk: hub.

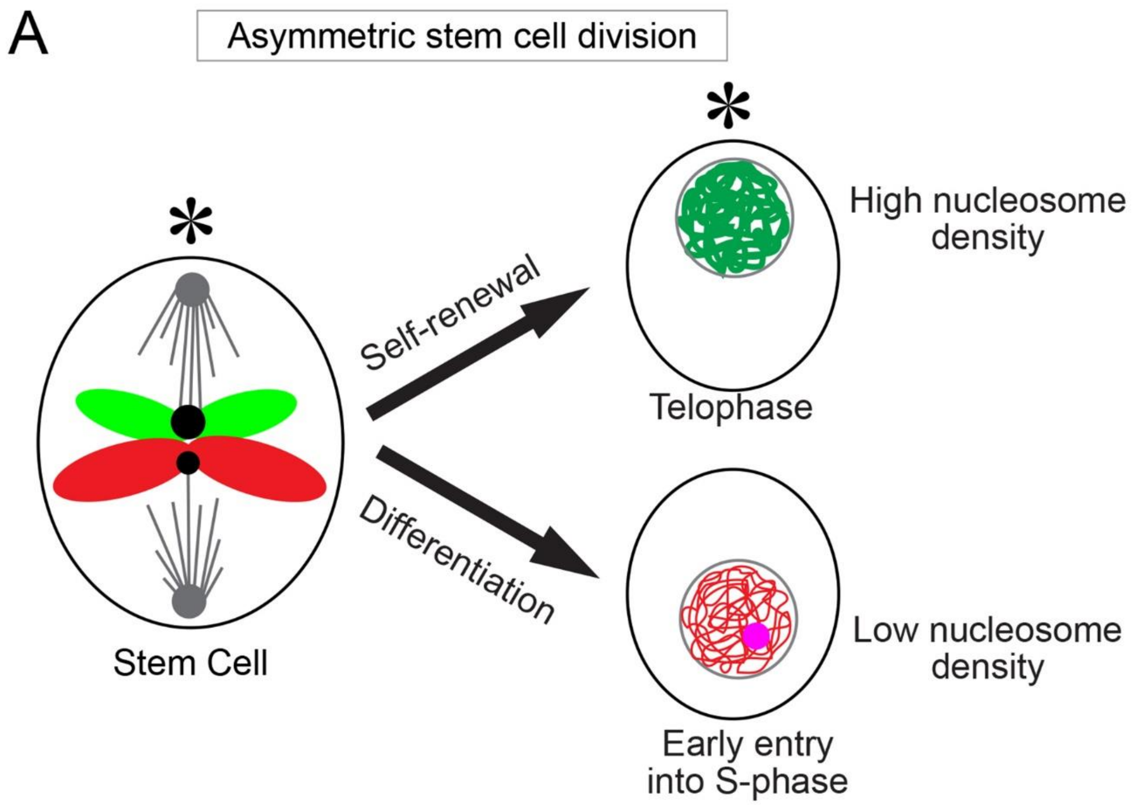

**Figure S4.** (**A**) Cartoon model of asymmetric entry into the next S-phase after asymmetric GSCs division.

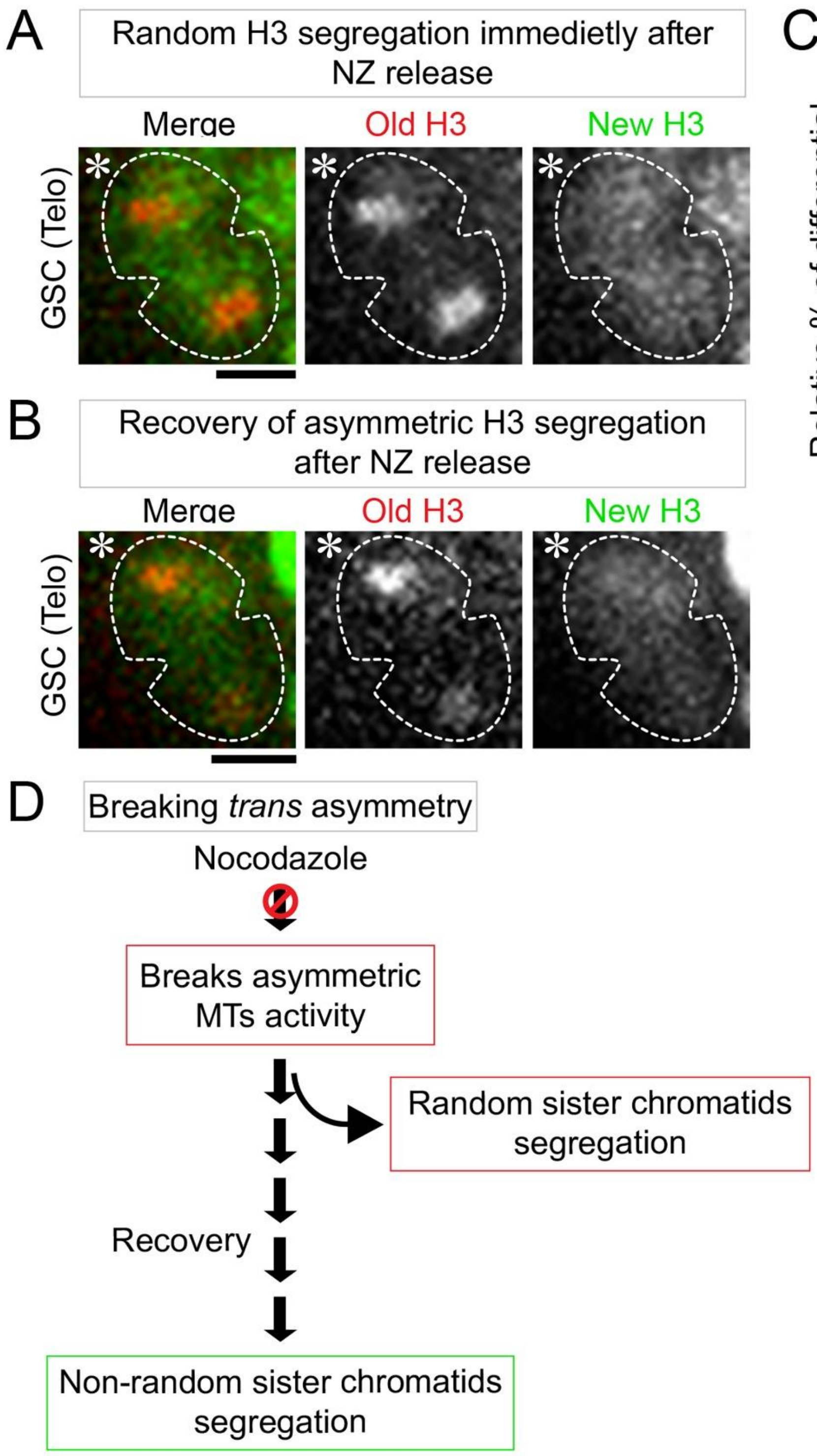

**Figure S5.** (**A**) Live cell image of symmetric H3 distribution immediately following release from NZ. (**B**) Live cell image after recovery from NZ showing asymmetric old histone. (**C**)

Calculation of the fraction of GSCs with asymmetric recovery using a cutoff for calling asymmetry compaction factor as described in the methods. (**D**) Model of randomize and recovery after NZ addition. Scale bars are 5 μm. Asterisk: hub.

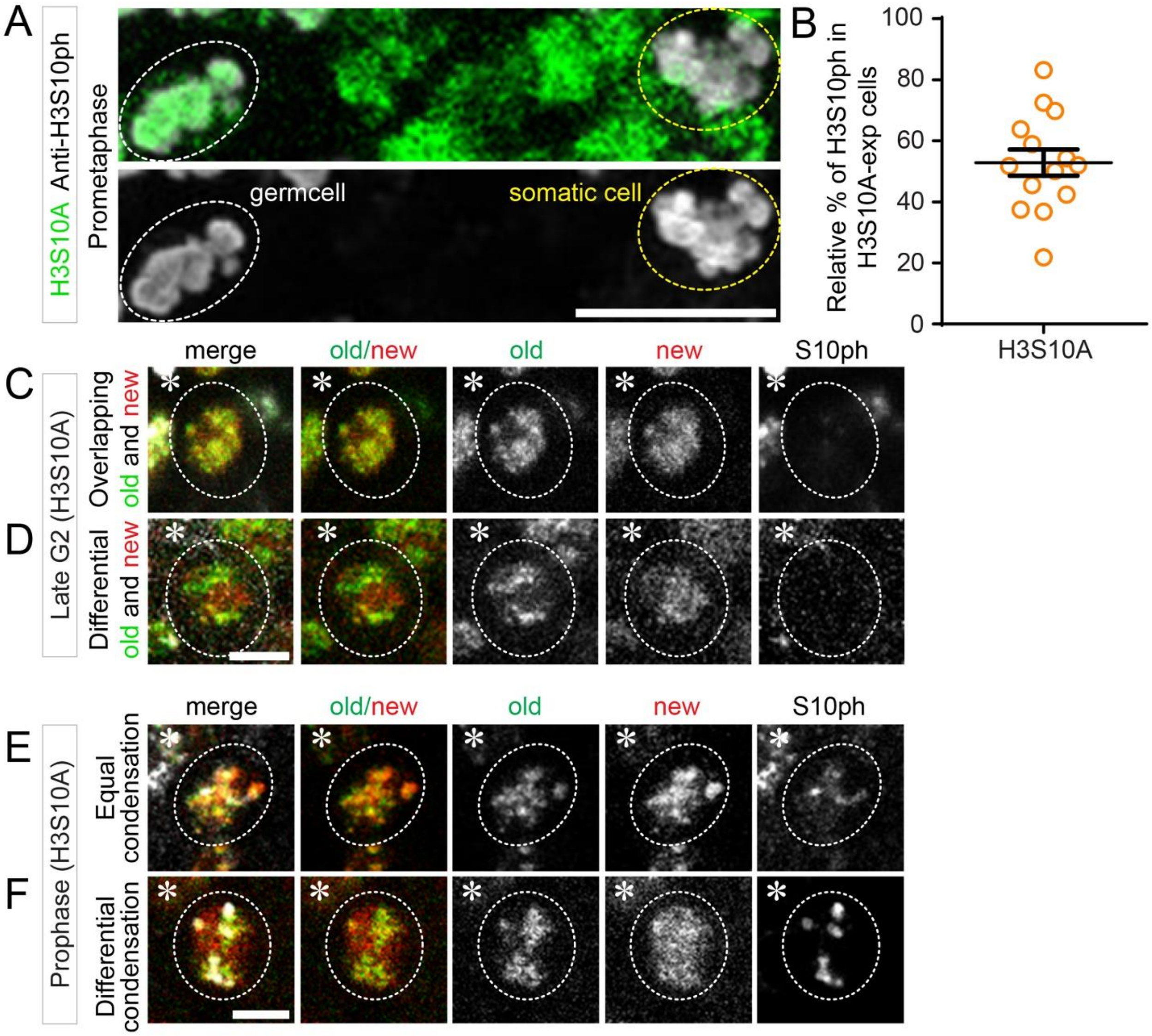

**Figure S6.** (**A**) Image of an adjacent mitotic somatic and germ cell in the H3S10A background showing that the level of H3S10ph is reduced in the germ cell which expresses H3S10A. (**B**) Quantification of the relative levels of H3S10ph between H3S10A germ cells and adjacent wild type somatic cells shows the reduction in H3S10ph levels in the mutant [average = 52.88 ± 4.31 (n=14)]. (**C**, **D**) H3S10A results in heterogeneous separation patterns of old and new histones in late G2, showing both overlapping (**C**) and differential (**D**) separation patterns. (**E**, **F**) H3S10A results in heterogeneous condensation patterns of old and new histones in prophase as well, showing both equal (**E**) and differential (**F**) condensation patterns. Scale bars are 10 μm (**A**) 2 μm (**C-F**). Asterisk: hub. All ratios = average ± SE.

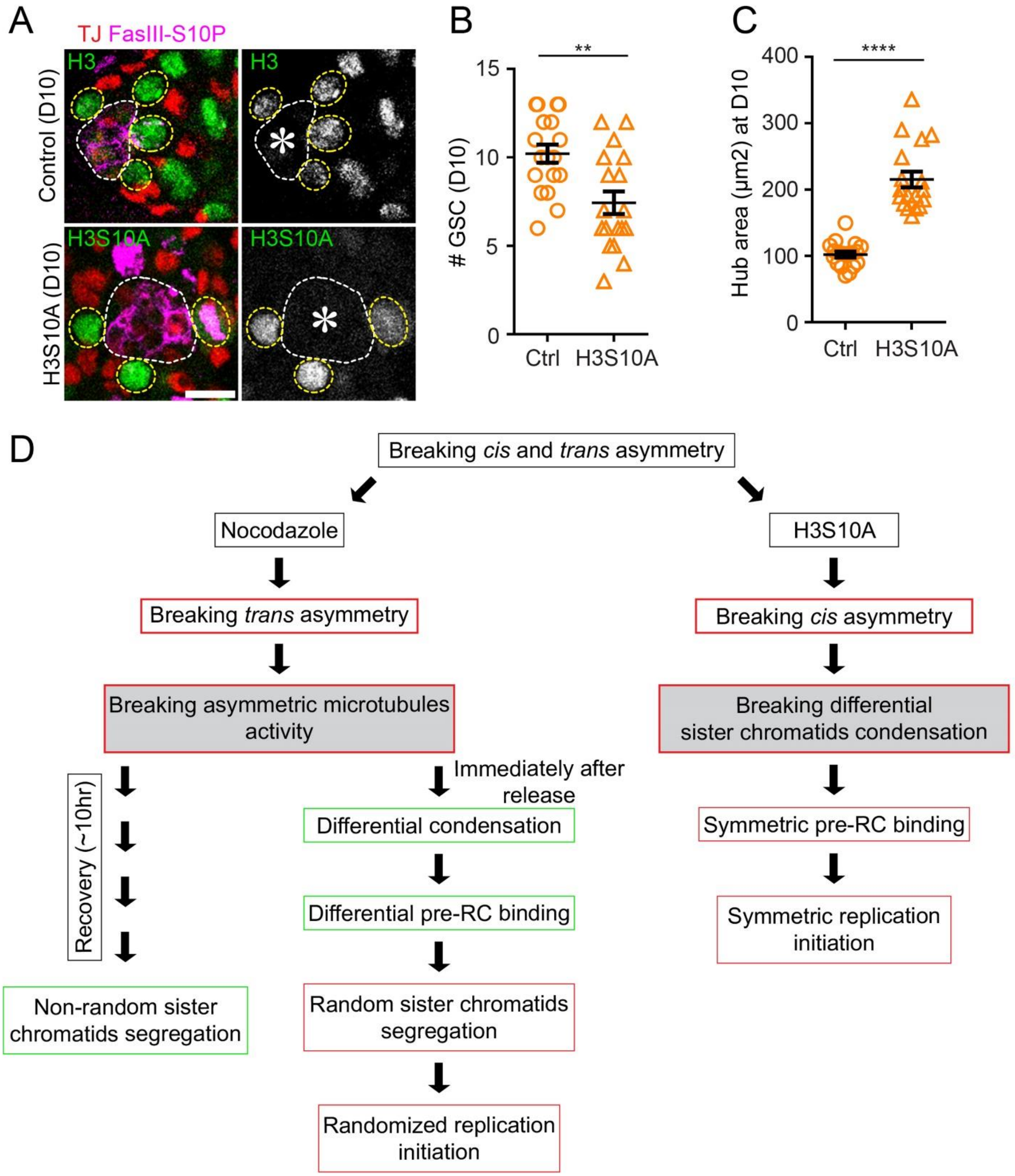

**Figure S7.** (**A**) Images showing a decreased number of GSCs by 10 days after eclosion in H3S10A compared to wild type and hub expansion in the mutant. (**B**) Quantification showing the reduced number of GSCs after 10 days post-eclosion [Ctrl, average = 10.22 ± 0.51 (n= 18);

H3S10A, average = 7.44 ± 0.64 (n=18)]. (**C**) Quantification of hub size after 10 days post-eclosion showing the expanded hub phenotype in the H3S10A mutant compared to wild type [Ctrl, average = 102.29 ± 4.49 (n=18) ; H3S10A, average = 215.41 ± 11.80 (n=18)]. (**D**) Comparison of the logical reasons why NZ results in randomized entry into the next S-phase while H3S10A induces symmetric entry into the next S-phase. Scale bars are 5 µm. Asterisk: hub. All ratios = average ± SE.

**Supplementary movie legends:**

**Movie S1. Related to Figure 1: Histone H3 nucleosome density asymmetry on sister chromatids during asymmetric GSC division.** Live cell imaging of the male *Drosophila* GSC expressing H3-mCherry. This video is a 3D reconstruction of individual images (10 × 1-µm interval optical sections per frame). ACD showing a high level of H3 nucleosome on sister chromatids segregated in GSCs in telophase. The video was acquired at 5-min intervals for 20-30 minutes. Metaphase is used as a landmark to define time point zero, and other time points are labeled as minus minutes prior to metaphase. The quantification is shown in Figure 1. Asterisk: hub. Scale bar: 5µm.

**Movie S2. Related to Figure 1: Symmetric H3 nucleosome density on sister chromatids during symmetric SG division.** Live cell imaging of the male *Drosophila* SG expressing H3-mCherry. This video is a 3D reconstruction of individual images (10 × 1-µm interval optical sections per frame). Symmetric cell division showing an equal level of H3 nucleosome on sister chromatids segregated in daughter cells in telophase. The video was acquired at 5-min intervals for 20-30 minutes. Metaphase is used as a landmark to define time point zero, and other time points are labeled as minus minutes prior to metaphase. The quantification is shown in Figure 1. Asterisk: hub. Scale bar: 5µm.

**Movie S3. Related to Figure 1: Histone H4 nucleosome density asymmetry on sister chromatids during asymmetric GSC division.** Live cell imaging of the male *Drosophila* GSC expressing H4-mCherry. This video is a 3D reconstruction of individual images (10 × 1-µm interval optical sections per frame). ACD showing a high level of H4 nucleosome on sister chromatids segregated in GSCs in telophase. The video was acquired at 5-min intervals for 20-30 minutes. Metaphase is used as a landmark to define time point zero, and other time points are labeled as minus minutes prior to metaphase. The quantification is shown in Figure 1. Asterisk: hub. Scale bar: 5µm.

**Movie S4. Related to Figure 1: Symmetric H4 nucleosome density on sister chromatids during symmetric SG division.** Live cell imaging of the male *Drosophila* SG expressing H4-mCherry. This video is a 3D reconstruction of individual images (10 × 1-µm interval optical sections per frame). Symmetric cell division showing an equal level of H4 nucleosome on sister chromatids segregated in daughter cells in telophase. The video was acquired at 5-min intervals for 20-30 minutes. Metaphase is used as a landmark to define time point zero, and other time points are labeled as minus minutes prior to metaphase. The quantification is shown in Figure 1. Asterisk: hub. Scale bar: 5µm.

**Movie S5. Related to Figure 1: Histone H2A nucleosome density asymmetry on sister chromatids during asymmetric GSC division.** Live cell imaging of the male *Drosophila* GSC expressing H2A-mCherry. This video is a 3D reconstruction of individual images (10 × 1-μm interval optical sections per frame). ACD showing a high level of H2A nucleosome on sister chromatids segregated in GSCs in telophase. The video was acquired at 5-min intervals for 20-30 minutes. Metaphase is used as a landmark to define time point zero, and other time points are labeled as minus minutes prior to metaphase. The quantification is shown in Figure 1. Asterisk: hub. Scale bar: 5μm.

**Movie S6. Related to Figure 1: Symmetric H2A nucleosome density on sister chromatids during symmetric SG division.** Live cell imaging of the male *Drosophila* SG expressing H2A-mCherry. This video is a 3D reconstruction of individual images (10 × 1-μm interval optical sections per frame). Symmetric cell division showing an equal level of H2A nucleosome on sister chromatids segregated in daughter cells in telophase. The video was acquired at 5-min intervals for 20-30 minutes. Metaphase is used as a landmark to define time point zero, and other time points are labeled as minus minutes prior to metaphase. The quantification is shown in Figure 1. Asterisk: hub. Scale bar: 5μm.

**Movie S7. Related to Figure 2: Condensation dynamics of old and new histone H3 enriched chromatin during asymmetric GSC division.** Live cell imaging of the dual-color histone, old H3-mCherry and new H3-EGFP, in a *Drosophila* male GSC. This video is a 3D reconstruction of individual images (10× 1-μm interval optical sections per frame), showing a differential condensation of old H3 chromatin and new H3 chromatin during mitosis. The video was acquired at 5-min intervals for 30-40 minutes to capture mitosis. Metaphase is used as a landmark to define time point zero, and other time points are labeled as minus minutes prior to metaphase. The quantification is shown in Figure 2. Asterisk: hub. Scale bar: 5μm.

**Movie S8. Related to Figure 2: Condensation dynamics of old and new histone H3 enriched chromatin during symmetric male *Drosophila* spermatogonial division.** Live cell imaging of the dual-color histone, old H3-mCherry and new H3-EGFP, in a *Drosophila* male GSC. This video is a 3D reconstruction of individual images (10 × 1-μm interval optical sections per frame), showing an equal condensation of old H3 chromatin and new H3 chromatin during mitosis. The video was acquired at 5-min intervals for 30-40 minutes to capture mitosis. Metaphase is used as a landmark to define time point zero, and other time points are labeled as minus minutes prior to metaphase. The quantification is shown in Figure 2. Asterisk: hub. Scale bar: 5μm.

**Supplementary tables and table legends:**

**Table S1. Related to Figure 1: H3 nucleosome density quantification.** H3 nucleosome density in telophase was quantified using high-resolution live-cell imaging of GSCs or SGs expressing H3-mCherry. Quantification shows the asymmetric nucleosome density on sister chromatids. The number represents the number of GSCs or SGs.

| | H3 | H3 |
|---|---|---|
| # | GSC/GB | SG1/SG2 |
| 1 | 1.79652 | 0.99816 |
| 2 | 1.87503 | 1.03214 |
| 3 | 1.24935 | 0.8575 |
| 4 | 1.23579 | 1.46861 |
| 5 | 1.2477 | 1.41326 |
| 6 | 1.39529 | 1.10108 |
| 7 | 1.81993 | 1.49463 |
| 8 | 1.41707 | 1.33178 |
| 9 | 1.50669 | 1.24479 |
| 10 | 1.35875 | 1.13987 |
| 11 | 1.63876 | 1.13003 |
| 12 | 1.48409 | 0.83183 |
| 13 | | 1.0212 |
| 14 | | 0.76469 |
| 15 | | 0.68237 |

**Table S2. Related to Figure 1: H4 nucleosome density quantification.** H4 nucleosome density in telophase was quantified using high-resolution live-cell imaging of GSCs or SGs expressing H4-mCherry. Quantification shows the asymmetric nucleosome density on sister chromatids. The number represents the number of GSCs or SGs.

| | H4 | H4 |
|---|---|---|
| # | GSC/GB | SG1/SG2 |
| 1 | 1.65121 | 0.78266 |
| 2 | 1.41131 | 0.87647 |
| 3 | 1.8106 | 1.05908 |
| 4 | 1.37548 | 0.96316 |
| 5 | 1.48215 | 0.9148 |
| 6 | 1.45867 | 0.81164 |
| 7 | 1.24271 | 1.40292 |
| 8 | 1.24297 | 1.07874 |
| 9 | 1.6608 | 0.99753 |
| 10 | 1.18161 | 1.11585 |
| 11 | 1.23359 | 1.02532 |
| 12 | 1.42363 | 0.86985 |
| 13 | 1.2304 | 1.04652 |

| | | |
|---|---|---|
| 14 | | 1.32634 |

**Table S3. Related to Figure S1: H2A nucleosome density quantification.** H2A nucleosome density in telophase was quantified using high-resolution live-cell imaging of GSCs or SGs expressing H2A-mCherry. Quantification shows the asymmetric nucleosome density on sister chromatids. The number represents the number of GSCs or SGs.

| | H2A | H2A |
|---|---|---|
| # | GSC/GB | SG1/SG2 |
| 1 | 1.713317 | 1.021659 |
| 2 | 1.507668 | 0.871567 |
| 3 | 1.216831 | 0.959251 |
| 4 | 1.23095 | 0.845858 |
| 5 | 1.303911 | 1.111325 |
| 6 | 1.221695 | 0.855221 |
| 7 | 1.239301 | 1.24233 |
| 8 | 1.244802 | 1.080725 |
| 9 | 1.398382 | 0.711259 |
| 10 | 1.389339 | 1.105897 |
| 11 | 1.354794 | 1.304479 |
| 12 | 1.128054 | 1.065765 |
| 13 | 1.235888 | 1.37247 |
| 14 | 1.40138 | 0.805808 |
| 15 | 1.590183 | 1.191536 |
| 16 | 1.151875 | 1.136097 |
| 17 | 1.311355 | 0.922961 |
| 18 | | 1.096929 |
| 19 | | 1.349674 |
| 20 | | 1.046363 |
| 21 | | 0.942935 |
| 22 | | 1.351117 |
| 23 | | 0.935996 |
| 24 | | 1.058753 |
| 25 | | 1.10034 |

**Table S4. Related to Figure 1: Nucleosome density quantification.** H3 nucleosome density in S-phase was quantified using super-resolution imaging of chromatin fiber from early germcells expressing H3-eGFP. Quantification shows the asymmetric nucleosome density on sister chromatin fiber. The number represents the number of chromatin fibers.

| # | Log2(Leading/Lagging H3) |
|---|---|
| 1 | 1.083108 |
| 2 | 0.747224 |
| 3 | 0.245465 |
| 4 | -0.2996 |
| 5 | 0.144415 |
| 6 | -0.12454 |
| 7 | 1.462192 |
| 8 | 0.882492 |
| 9 | 1.900402 |
| 10 | -0.08197 |
| 11 | -0.39285 |
| 12 | 0.355208 |
| 13 | 0.675815 |
| 14 | 0.534285 |
| 15 | -0.23673 |
| 16 | -0.43068 |
| 17 | -0.7049 |
| 18 | 1.228543 |
| 19 | -0.52004 |
| 20 | 1.019822 |
| 21 | 1.826114 |
| 22 | 0.462984 |
| 23 | 0.402327 |
| 24 | 1.147252 |
| 25 | -0.0108 |
| 26 | 1.391923 |
| 27 | 0.703404 |
| 28 | 1.744645 |
| 29 | 0.135108 |

**Table S5. Related to Figure 2: Quantification of sister chromatids condensation.** Used dual-color histone labeling system (UASp-FRT-histone-mCherry-PolyA-FRT-histone-eGFP-PolyA transgene) to distinguish old and new H3 enriched chromatin. Prometaphase GSCs or SGs of high-resolution live-cell images were used for quantification. Late-prophase and prometaphase of fixed cell images were used for quantification. Quantification shows the differential-sister chromatids condensation in *Drosophila* male GSCs but not in SGs. The number represents the number of GSCs or SGs.

| | Live cell imaging | | Fixed cell imaging | |
|---|---|---|---|---|
| # | GSC | SG | GSC | SG |
| 1 | 4.029748284 | 1.2586207 | 2.071428571 | 1.4963119 |
| 2 | 3.09418146 | 0.9894737 | 3.858356941 | 1.1568627 |
| 3 | 5.728835979 | 1.3882353 | 2.099150142 | 1.578271 |
| 4 | 4.043743642 | 1.1226415 | 1.953007519 | 1.4683841 |
| 5 | 5.828125 | 0.9915966 | 3.006232687 | 1.4208494 |
| 6 | 4.967651195 | 1.0350877 | 1.943894389 | 1.4790875 |
| 7 | 6.192176871 | 1.2461538 | 2.810741688 | 1.1606061 |
| 8 | 4.174358974 | 1.1666667 | 2.243697479 | 1.09375 |
| 9 | 3.532580365 | 1.3914894 | 3.626943005 | 1.4291188 |
| 10 | 4.142857143 | 0.8438538 | 1.601073345 | 1.2916667 |
| 11 | 3.373633441 | 1.3670412 | 2.594972067 | 1.2924528 |
| 12 | | 1.0462287 | 6.96485623 | 1.1497711 |
| 13 | | 2.062201 | 1.597202797 | |
| 14 | | 1.0178571 | 1.778645833 | |
| 15 | | 2.3282828 | 1.471317829 | |
| 16 | | 1.1282051 | | |
| 17 | | 1.5113636 | | |
| 18 | | 1.8988764 | | |

**Table S6. Related to Figure 2: Differential colocalization of histone H3 Serine 10 phosphorylation with old and new H3 enriched chromatin.**

| # | H3S10ph with Old H3 | H3S10ph with New H3 |
|---|---|---|
| 1 | 0.4260082 | 0.08524515 |
| 2 | 0.3450814 | 0.03980097 |
| 3 | 0.3088183 | 0.06380519 |
| 4 | 0.4472051 | 0.50134893 |
| 5 | 0.4944739 | 0.09393801 |
| 6 | 0.5046727 | 0.40986226 |
| 7 | 0.5796574 | 0.38435161 |
| 8 | 0.3948776 | 0.15332015 |
| S9 | 0.280832 | 0.12677746 |
| 10 | 0.75566 | 0.1882747 |
| 11 | 0.7901022 | -0.0802569 |

| | | |
|---|---|---|
| 12 | 0.8340661 | -0.01767 |
| 13 | 0.4077583 | -0.0372955 |
| 14 | 0.3746972 | 0.62775392 |
| 15 | 0.4393611 | 0.58747564 |

**Table S7. Related to Figure 3: Cdc6 segregation during asymmetric GSCs division and symmetric SG division using super-resolution live snapshot technique.** Cdc6 was tagged endogenously with mCherry (cdc6-mCherry) using the CRISPR/Cas9 technique and used for the live imaging.

| # | GB/GSC | SG1/SG2 |
|---|---|---|
| 1 | 1.9252475 | 1.27117 |
| 2 | 1.7367384 | 0.95729 |
| 3 | 1.8713297 | 1.14654 |
| 4 | 1.8863495 | 1.08313 |
| 5 | 2.2302022 | 1.04434 |
| 6 | 1.4254525 | 1.54837 |
| 7 | 1.8953589 | 0.91575 |
| 8 | 2.8933531 | 1.11509 |
| 9 | 1.5137818 | 1.06324 |
| 10 | 1.2706748 | 1.16787 |
| 11 | 1.4340685 | 1.06705 |
| 12 | 1.1385365 | 1.31529 |
| 13 | 1.4317549 | 1.19664 |
| 14 | 1.7693503 | 1.26831 |
| 15 | | 1.17556 |

**Table S8. Related to Figure 4: Asymmetric DNA replication initiation as shown by PCNA.**

| # | Log2 (GSC/GB PCNA) | Log2(SG1/SG2 PCNA) |
|---|---|---|
| 1 | -1.402563387 | 0.012740987 |
| 2 | -1.857980995 | -0.031042422 |
| 3 | -2.502500341 | -0.113743366 |
| 4 | -0.887525271 | -0.041928834 |
| 5 | 0.029146346 | 0.218806341 |
| 6 | -0.343094907 | 0.132207909 |
| 7 | 0.527349772 | -0.009847786 |
| 8 | 0.064937942 | -0.02529402 |
| 9 | -0.099130053 | -0.065095028 |
| 10 | -0.581265553 | -0.206450877 |
| 11 | -0.807354922 | -0.05703179 |
| 12 | 0.277533976 | 0.017620373 |
| 13 | 0.065621994 | -0.048196686 |

| | | |
|---|---|---|
| 14 | 0.63005039 | 0.174725988 |
| 15 | -0.2296055 | -0.122860929 |
| 16 | -0.152003093 | 0.032498335 |
| 17 | 0.031042422 | 0.042973782 |
| 18 | -0.062914736 | -0.188761643 |
| 19 | -1.459431619 | -0.039649077 |
| 20 | -0.319459839 | -0.139162748 |
| 21 | 0 | 0.259212695 |
| 22 | 0.443950191 | -0.006617879 |
| 23 | 0.339486466 | 0.13492958 |
| 24 | -1.415037499 | 0.415037499 |
| 25 | 1.123988717 | -0.434262067 |
| 26 | 0.227068909 | 0.075442556 |
| 27 | -0.083768358 | 0.103093493 |
| 28 | 0.46712601 | 0.263962482 |
| 29 | 0.199769512 | 0.099535674 |
| 30 | 0.509013647 | -0.063586683 |
| 31 | 0.030373649 | 0.216133431 |
| 32 | -0.373458396 | -0.151722667 |
| 33 | 0.27064759 | |

**Table S9. Related to Figure 4: Asymmetric DNA replication initiation as shown by EdU.**

| # | Log2(GB/GSC EdU) | Log2(SG1/SG2 EdU) |
|---|---|---|
| 1 | -1.05154612 | 0.012740987 |
| 2 | -1.596935142 | -0.031042422 |
| 3 | -3 | -0.113743366 |
| 4 | -1.944858446 | -0.11321061 |
| 5 | -3.387023123 | -0.05703179 |
| 6 | -0.137503524 | 0.017620373 |
| 7 | 0.029146346 | -0.048196686 |
| 8 | 0.067080121 | 0.174725988 |
| 9 | -0.124328135 | -0.122860929 |
| 10 | -0.070389328 | 0.032498335 |
| 11 | -1.485426827 | -0.052614626 |
| 12 | -0.378511623 | -0.011097709 |
| 13 | -0.564498398 | 0.080170349 |
| 14 | -0.701965111 | 0.192645078 |
| 15 | -0.95419631 | -0.066769012 |
| 16 | -0.797507136 | -0.321928095 |
| 17 | -1.073406727 | 0.415037499 |
| 18 | -2.123525746 | 0.211504105 |
| 19 | -0.433040734 | -0.205318908 |
| 20 | -0.7589919 | 0.072900547 |

| | | |
|---|---|---|
| 21 | -1.485426827 | 0.009073584 |
| 22 | 0.167456746 | 0.128866595 |
| 23 | -1.485426827 | 0.034351505 |
| 24 | -0.360175564 | 0.131672633 |
| 25 | -0.132755209 | -0.150561135 |
| 26 | -0.022026306 | |
| 27 | -0.042644337 | |
| 28 | -0.474626296 | |
| 29 | 0.604787479 | |
| 30 | 0 | |
| 31 | -0.10433666 | |
| 32 | 0.705386877 | |
| 33 | 0 | |
| 34 | 0.457850583 | |
| 35 | 0 | |
| 36 | -0.171611378 | |
| 37 | 0.167456746 | |

**Table S10. Related to Figure 5: Histone H3 segregation pattern immediately after nocodazole release and after ~10hrs recovery.**

| | NZ release | | NZ Recovery | |
|---|---|---|---|---|
| | Old H3 | New H3 | Old H3 | New H3 |
| # | GSC/GB | GB/GSC | GSC/GB | GB/GSC |
| 1 | 1.482336224 | 0.77609528 | 1.778144954 | 0.993901749 |
| 2 | 0.855046833 | 1.039199333 | 1.66260378 | 0.635398958 |
| 3 | 0.944940051 | 1.02671019 | 1.505044843 | 0.904210526 |
| 4 | 1.11008235 | 0.921893779 | 2.385225259 | 0.744099379 |
| 5 | 0.671087782 | 1.452010706 | 1.038143567 | 1.187762997 |
| 6 | 0.901072346 | 1.155459329 | 1.675295814 | 0.83521727 |
| 7 | 1.093413811 | 0.72857707 | 1.633093817 | 0.707618406 |
| 8 | 0.889588121 | 1.145887571 | 1.070594595 | 1.044318656 |
| 9 | 1.057759221 | 0.805306184 | 1.872997446 | 0.666752354 |
| 10 | 0.782211152 | 0.942145172 | 1.05179188 | 0.941362019 |
| 11 | 1.255861906 | 0.982216817 | 1.405161523 | 0.712148467 |
| 12 | 0.86744603 | 1.41846703 | 1.538197078 | 1.312581816 |
| 13 | 1.289635091 | 0.66895723 | 1.44014457 | 0.684998957 |
| 14 | 0.803190272 | 0.930187232 | 1.250846911 | 0.813995824 |
| 15 | 1.138100083 | 0.881104552 | 1.098650105 | 1.075498158 |
| 16 | 0.883464329 | 1.381390246 | 2.446066746 | 0.43324518 |
| 17 | 1.324686941 | 0.893723426 | 1.412264906 | 0.924281653 |
| 18 | 1.10417312 | 1.00980101 | 1.474716981 | 0.721516298 |
| 19 | 0.9696303 | 0.894523759 | 1.321547872 | 1.02843952 |
| 20 | 1.647926736 | 1.04764797 | 1.033378739 | 0.997320778 |

| | | |
|---|---|---|
| 21 | 1.854211663 | 0.605085603 |
| 22 | 0.998369392 | 1.03658257 |
| 23 | 0.932589688 | 0.9537048 |
| 24 | 1.446077574 | 0.916642484 |
| 25 | 0.85851039 | 0.989764021 |
| 26 | 0.875320395 | 1.067987756 |

**Table S11. Related to Figure 5: Old and new histone H3 enriched sister chromatids condense differentially after nocodazole arrest and release in male *Drosophila* GSCs.**

| # | Compaction factor (GSC) |
|---|---|
| 1 | 1.599206349 |
| 2 | 1.727427598 |
| 3 | 1.715746421 |
| 4 | 1.666666667 |
| 5 | 3.621052632 |
| 6 | 2.667706708 |
| 7 | 1.748120301 |
| 8 | 1.754330709 |
| 9 | 1.223021583 |
| 10 | 2.54601227 |
| 11 | 3.996336996 |
| 12 | 1.69212963 |
| 13 | 4.385826772 |
| 14 | 4.975177305 |
| 15 | 3.269430052 |
| 16 | 1.258751903 |

**Table S12. Related to Figure 5: Cdc6 segregation randomized after nocodazole arrest and release in male *Drosophila* GSCs.** Super-resolution live snapshots technique was used for imaging telophase GSCs expressing endogenously tagged cdc6-mCherry.

| # | GB/GSC |
|---|---|
| 1 | 1.10409 |
| 2 | 0.85016 |
| 3 | 0.61835 |
| 4 | 1.08103 |
| 5 | 2.45921 |
| 6 | 0.72114 |
| 7 | 1.74348 |

| | |
|---|---|
| 8 | 0.49415 |
| 9 | 0.4352 |
| 10 | 0.54374 |
| 11 | 1.55773 |
| 12 | 0.99717 |
| 13 | 1.06662 |

**Table S13. Related to Figure 5: Quantification of PCNA after nocodazole arrest and release in male *Drosophila* GSCs.**

| # | Corrected Sum of Area | Log2(GSC/GB PCNA) |
|---|---|---|
| 1 | 23.7592 | 3.540163 |
| 2 | 34.7888 | 7.738451 |
| 3 | 67.9704 | 0.27838 |
| 4 | 63.0536 | 0.498076 |
| 5 | 99.0384 | 0.518708 |
| 6 | 87.696 | 0.893394 |
| 7 | 75.564 | -0.42938 |
| 8 | 35.7016 | -6.06874 |
| 9 | 48.9048 | -0.97633 |
| 10 | 43.2328 | -1.10238 |
| 11 | 33.1816 | 0.113075 |
| 12 | 19.4736 | -0.82594 |
| 13 | 40.5224 | -2.01027 |
| 14 | 39.4528 | 2.086565 |
| 15 | 43.1728 | 1.42278 |
| 16 | 51.8184 | -1.02341 |
| 17 | 57.4732 | -1.69432 |
| 18 | 47.9468 | -0.19168 |
| 19 | 43.0476 | -0.31009 |
| 20 | 39.1044 | -0.39259 |
| 21 | 57.5556 | 0.462189 |
| 22 | 50.4808 | -0.36343 |
| 23 | 51.6456 | -1.79623 |
| 24 | 60.9416 | -0.38361 |
| 25 | 31.8256 | -1.06247 |
| 26 | 45.284 | 0.148678 |
| 27 | 13.1088 | 0.392048 |
| 28 | 15.442 | -1.76504 |

**Table S14. Related to Figure 6: Quantification of sister chromatids condensation in histone H3S10A expressing GSCs.**

| # | Compaction factor (GSCs) |
|---|---|
| 1 | 1.4754335 |
| 2 | 1.1706865 |
| 3 | 1.4520202 |
| 4 | 1.3629403 |
| 5 | 1.9347826 |
| 6 | 1.0537897 |
| 7 | 1.4804965 |
| 8 | 1.2882096 |
| 9 | 1.2436647 |
| 10 | 1.6101695 |
| 11 | 1.7807018 |
| 12 | 1.5437158 |
| 13 | 1.69 |
| 14 | 1.5506173 |
| 15 | 1.1711538 |
| 16 | 0.993763 |
| 17 | 1.9057143 |
| 18 | 1.1404762 |
| 19 | 1.3142857 |
| 20 | 1.0862069 |
| 21 | 1.5686567 |
| 22 | 1.1397206 |
| 23 | 1.2682119 |
| 24 | 1.9630314 |
| 25 | 1.1618887 |
| 26 | 1.1747967 |
| 27 | 1.5056818 |
| 28 | 1.1832718 |
| 29 | 1.4859438 |
| 30 | 1.1963928 |
| 31 | 1.4931507 |
| 32 | 1.4096386 |
| 33 | 1.8179104 |
| 34 | 1.1699346 |
| 35 | 1.2293233 |
| 36 | 1.5666667 |
| 37 | 1.510574 |
| 38 | 1.2033898 |
| 39 | 1.283293 |
| 40 | 1.3443709 |
| 41 | 0.8649007 |
| 42 | 1.040678 |
| 43 | 0.840708 |
| 44 | 1.1876523 |

| | |
|---|---|
| 45 | 1.39762 |

**Table S15. Related to Figure 6: Quantification of old and new H3 segregation in H3S10A expressing GSCs.**

| # | Old H3 | New H3 |
|---|---|---|
| 1 | 0.922 | 0.94 |
| 2 | 0.81 | 0.79 |
| 3 | 0.95 | 0.851 |
| 4 | 1.12 | 0.97 |
| 5 | 1.03 | 1.21 |
| 6 | 0.79 | 1.05 |
| 7 | 1.08 | 1.3 |
| 8 | 0.98 | 0.8 |
| 9 | 0.66 | 0.7 |
| 10 | 0.925 | 1.07 |
| 11 | 1.012 | 1.091 |
| 12 | 1.03 | 0.952 |
| 13 | 0.93 | 1.18 |
| 14 | 1.027 | 0.93 |

**Table S16. Related to Figure 7: Quantification of replication initiation using EdU staining in H3S10A expressing GSCs.**

| # | Sum of Area Corrected | Log2(EdU ratio [GSC/GB]) |
|---|---|---|
| 1 | 40.8996 | -0.1082 |
| 2 | 34.5247 | 0.06464 |
| 3 | 48.5093 | 0.36099 |
| 4 | 39.3558 | -0.3701 |
| 5 | 25.6071 | -0.172 |
| 6 | 27.9496 | -0.549 |
| 7 | 25.1545 | -0.3725 |
| 8 | 46.2749 | -0.2957 |
| 9 | 49.7259 | -0.2172 |
| 10 | 45.5669 | -0.0114 |
| 11 | 35.4683 | 0.07053 |
| 12 | 43.1515 | 0.43736 |
| 13 | 72.7349 | -0.0531 |
| 14 | 48.586 | 0.05175 |
| 15 | 22.8015 | -0.2993 |
| 16 | 53.0408 | 0.29043 |
| 17 | 37.8298 | 0.01756 |
| 18 | 47.7638 | -0.0533 |

| | | |
|---|---|---|
| 19 | 42.8967 | -0.2485 |
| 20 | 44.4598 | -0.5662 |
| 21 | 40.046 | -0.0849 |
| 22 | 34.0715 | -0.0924 |
| 23 | 23.1371 | -0.1636 |
| 24 | 45.0219 | 0.01412 |
| 25 | 50.3983 | -0.0165 |

**Supplemental References**

Blythe, S.A., and Wieschaus, E.F. (2016). Establishment and maintenance of heritable chromatin structure during early Drosophila embryogenesis. Elife *5*.

Hime, G.R., Brill, J.A., and Fuller, M.T. (1996). Assembly of ring canals in the male germ line from structural components of the contractile ring. J Cell Sci *109 ( Pt 12)*, 2779-2788.

Maddox, P.S., Portier, N., Desai, A., and Oegema, K. (2006). Molecular analysis of mitotic chromosome condensation using a quantitative time-resolved fluorescence microscopy assay. Proc Natl Acad Sci U S A *103*, 15097-15102.

McCleland, M.L., Shermoen, A.W., and O'Farrell, P.H. (2009). DNA replication times the cell cycle and contributes to the mid-blastula transition in Drosophila embryos. J Cell Biol *187*, 7-14.

Ranjan, R., and Chen, X. (2021). Super-Resolution Live Cell Imaging of Subcellular Structures. J Vis Exp.

Ranjan, R., Snedeker, J., and Chen, X. (2019). Asymmetric Centromeres Differentially Coordinate with Mitotic Machinery to Ensure Biased Sister Chromatid Segregation in Germline Stem Cells. Cell Stem Cell *25*, 666-681 e665.

Tran, V., Lim, C., Xie, J., and Chen, X. (2012). Asymmetric division of Drosophila male germline stem cell shows asymmetric histone distribution. Science *338*, 679-682.

Van Doren, M., Williamson, A.L., and Lehmann, R. (1998). Regulation of zygotic gene expression in Drosophila primordial germ cells. Curr Biol *8*, 243-246.

Wooten, M., Li, Y., Snedeker, J., Nizami, Z.F., Gall, J.G., and Chen, X. (2020). Superresolution imaging of chromatin fibers to visualize epigenetic information on replicative DNA. Nat Protoc *15*, 1188-1208.

Yamashita, Y.M., Jones, D.L., and Fuller, M.T. (2003). Orientation of asymmetric stem cell division by the APC tumor suppressor and centrosome. Science *301*, 1547-1550.

Yamashita, Y.M., Mahowald, A.P., Perlin, J.R., and Fuller, M.T. (2007). Asymmetric inheritance of mother versus daughter centrosome in stem cell division. Science *315*, 518-521.
